# Brain shuttle target expression levels vary by individual, not by brain region, disease, age, or sex

**DOI:** 10.64898/2026.05.21.726908

**Authors:** Ana R. Santa-Maria, Sanjid Shahriar, Vasanth Chandrasekhar, Emma Luteijn, Robin Horber, Samuel Hornstein, Sasha Tkachev, Tyler Levy, Ivan Gregoretti, Claire Simpson, Majd Ariss, Vilas Menon, Donald Ingber, James Gorman

**Affiliations:** Wyss Institute for Biologically Inspired Engineering at Harvard University, Boston, MA 02215, USA; Sidney Kimmel Medical College, Thomas Jefferson University Hospital, Philadelphia, PA 19107, USA; Cell Signaling Technology, Danvers, MA 01923, USA; Columbia University Irving Medical Center, New York, NY 10032, USA; Vascular Biology Program and Department of Surgery, Boston Children’s Hospital and Harvard Medical School, Boston, MA 02115, USA; Harvard John Paulson School of Engineering and Applied Sciences, Cambridge, MA 02134 USA

## Abstract

Therapeutics fused to brain shuttles that exploit endogenous receptor-mediated transport at the blood–brain barrier (BBB) offer a promising strategy to deliver large molecule drugs and biologics to the CNS. A fundamental but untested assumption underlying their clinical development is that their endothelial receptor targets are consistently expressed between individuals and between patient populations. Here, we analyzed gene and protein expression of eleven canonical brain shuttle targets in isolated human brain microvascular endothelial cells and brain microvessels from 11 large cohorts, using single-cell and single-nucleus transcriptomics and quantitative proteomics. Expression was remarkably stable between brain regions, sexes, ages, and normal health versus four major neurodegenerative conditions, Alzheimer’s disease, Parkinson’s disease, Huntington’s disease, and amyotrophic lateral sclerosis, with 612 of 631 comparisons (97%) showing no significant difference. Regional heterogeneity of brain shuttle targets previously reported in rodent models was not observed in human tissue, and disease states had minimal impact in all diseases examined. In striking contrast, target abundance differed consistently among individuals in every demographic and clinical group, for all eleven targets and all data modalities. These findings establish individual receptor abundance as a critical and previously uncharacterized variable for brain shuttle translational research, including clinical trial design and patient stratification.

## INTRODUCTION

Despite decades of research, the effective brain delivery of large molecule therapies for neurodegenerative and other central nervous system (CNS) disorders remains a formidable challenge ^1-3^. While large-molecule biologics such as monoclonal antibodies, oligonucleotides, enzymes, and gene therapies have shown therapeutic promise for CNS diseases, poor blood-brain barrier (BBB) penetration presents a significant challenge to their clinical development ^1, 3^. The BBB, formed primarily by brain microvascular endothelial cells (BMECs) in contact with pericytes and astrocytes, tightly regulates the entry of most macromolecules into the brain parenchyma, including large molecule drugs ^3, 4^.

To address this challenge, a class of biologically inspired engineering strategies has emerged that use binding moieties, increasingly referred to as brain shuttles, that bind to target proteins expressed on the luminal surface of BMECs ^1, 3^, hijacking endogenous vesicular trafficking pathways to carry therapeutic payloads across the brain endothelial cells and into the CNS. The most widely studied brain shuttle targets are receptors that mediate receptor-mediated transcytosis (RMT), a process in which receptor engagement triggers endocytosis, directional vesicular trafficking, and abluminal exocytosis. RMT targets include the transferrin receptor (TFR) ^5, 6^, low-density lipoprotein receptor (LDLR) ^7^, LRP1/LRP8 ^8, 9^, and insulin receptor (INSR) ^10^. Additional brain shuttle targets with distinct endogenous membrane biology include IGF-1R ^11^, the solute carriers SLC2A1/GLUT1 ^12^, and SLC3A2/CD98hc ^13^, glycosylphosphatidylinositol (GPI)-anchored proteins (GPI-AP) ALPL ^14^ and CA4 ^15^; and the phospholipid flippase TMEM30A ^16^.

Recent clinical milestones have demonstrated that brain shuttle strategies can achieve effective CNS delivery in humans: an anti-TfR enzyme fusion (pabinafusp alfa) achieved CNS exposure sufficient for Japanese regulatory approval in Hunter’s syndrome ^17, 18^, and the TfR-targeted anti-amyloid antibody trontinemab has advanced to global Phase III registration trials for Alzheimer’s disease (AD) based on its promising Phase II trial results ^19, 20^. Yet the clinical translation of these brain transport technologies has proceeded with a largely unexamined assumption: that the endothelial receptor targets on which they depend are sufficiently and homogeneously expressed at the BBB in the patient populations being treated. Notably, no brain shuttle clinical program to date has proposed an approach to measure transport target receptor expression in human brain tissue, nor included such a measure as a stratification variable or correlative endpoint, either in the Hunter’s syndrome programs targeting TfR ^21, 22^, nor in ongoing AD ^19^ and Parkinson’s disease (PD) trials ^23, 24^. Prior studies have reported altered expression of individual brain shuttle targets in neurodegenerative disease, including reduced LRP1 and GLUT1 in the AD cerebrovasculature ^25-27^, increased TfR in amyloid mouse models ^28^, and downregulated IGF1R in post-mortem AD brain ^29^. However, these analyses have been conducted in small, heterogeneous cohorts and have not systematically evaluated the dimension of variability most directly relevant to clinical outcome: inter-individual differences in receptor expression between patients. This gap is consequential: for brain shuttle strategies in which CNS drug uptake is contingent on the presence and abundance of a single endothelial receptor, substantial differences in target expression between individual patients could be a primary determinant of variability in brain drug delivery, target engagement, and clinical responses observed between individual patients in clinical trials. The extent of such inter-individual variability in brain shuttle target expression, versus differences in expression between groups defined by disease state, brain region, age, or sex, remains unknown.

Addressing this question requires a large-scale, systematic characterization of brain shuttle target expression in human brain tissue, a standard that existing studies have not met. Prior analyses have relied on animal models ^30, 31^, small human cohorts ^32^, or single brain regions ^31, 32^, providing insufficient resolution to disentangle the relative contributions of region, disease, demographics, and individual biology to variability in brain shuttle target expression. This knowledge gap is critically important for the design of trials in AD, PD, Huntington’s disease (HD), and amyotrophic lateral sclerosis (ALS), in which disease pathology is known to alter BBB function ^33, 34^ and could further alter brain shuttle target protein expression. Here, we address this gap by integrating data from 11 large human cohorts spanning single-nucleus RNA sequencing (snRNA-seq), single-cell RNA sequencing (scRNA-seq), and quantitative proteomics to systematically analyze variation in the BMEC expression of eleven canonical brain shuttle targets with diverse endogenous functions. Using rigorous statistical analysis, we test for significant differences in target expression between brain regions, healthy and neurodegenerative disease states, sexes, ages and, critically, between individuals.

## RESULTS

### Expression of brain shuttle target genes is conserved between brain regions and disease states

We first assessed the extent to which brain shuttle target expression varies between brain regions and neurodegenerative conditions. Using published human snRNA-seq datasets including multiple brain regions in control subjects versus patients with four major neurodegenerative diseases, AD ^35-37^, PD ^38^, HD ^39^, and ALS ^40^, we bioinformatically extracted BMEC clusters using known canonical BMEC transcripts. For each brain region and disease state, we computed aggregate transcript abundance profiles for genes encoding transmembrane (TM) proteins and GPI-APs (Supplementary Data File 1). Pairwise Spearman’s rank correlations of these aggregate profiles between all regions and disease states were uniformly high, with coefficients ranging from 0.83 to 1.00 (Fig. 1), indicating that the global BMEC surface protein transcriptome is highly conserved between brain regions and largely preserved between these neurodegenerative disease states.

**Fig. 1:**
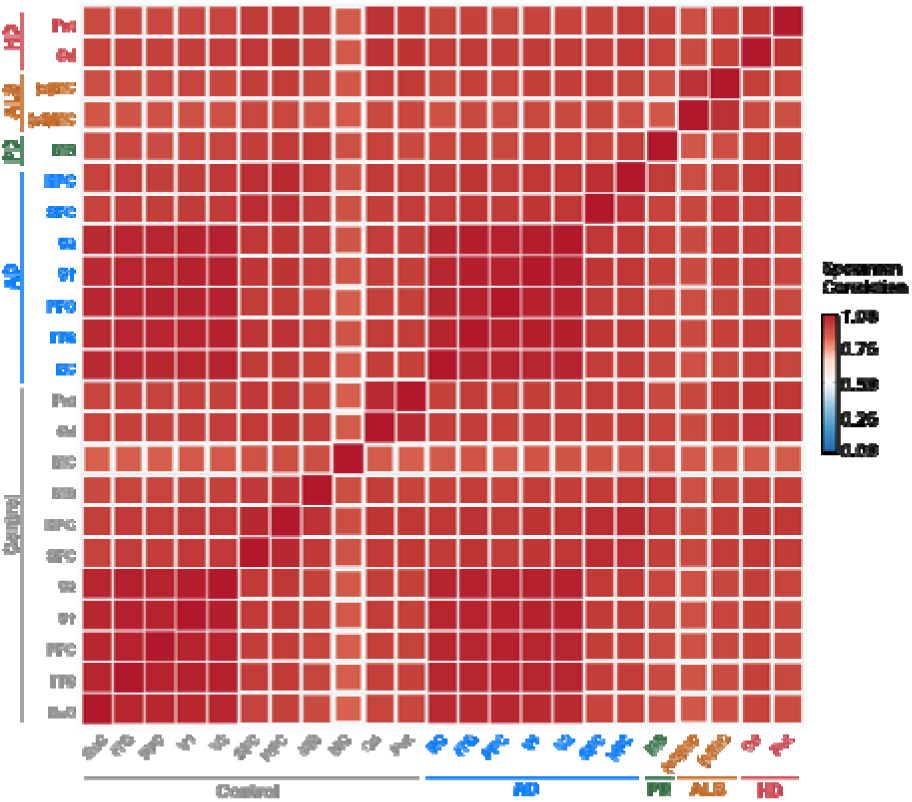
Transcriptomic profiles of genes encoding endothelial cell surface brain shuttle target proteins are conserved between brain regions and neurodegenerative diseases. The graphic shows Spearman’s correlations of aggregate expression (counts per million, CPM) derived from snRNA-seq datasets for genes encoding TM and GPI-anchored surface proteins in human BMECs. Expression data for multiple individuals were aggregated by brain region and disease state, including healthy controls and individuals with Alzheimer’s disease (AD), Parkinson’s disease (PD), Huntington’s disease (HD), and amyotrophic lateral sclerosis (ALS). Spearman’s rank correlation coefficients (ρ) ranged from 0.83 to 1.00, with square size proportional to correlation strength. Brain region abbreviations: EnC, entorhinal cortex; ITG, inferior temporal gyrus; PFC, prefrontal cortex; V1, visual cortex 1; V2, visual cortex 2; HPC, hippocampus; SFC, superior frontal cortex; MB, midbrain; MC, motor cortex; Cd, caudate; Put, putamen. Disease subtype abbreviations: c9MC, C9orf72 familial ALS; sMC, sporadic ALS.

### Brain shuttle target expression exhibits few significant differences between regions, diseases, sexes, and ages

To quantify how brain shuttle target expression differs in relation to key biological factors, we systematically compared gene expression levels between brain regions, disease states, sex, and age groups. Across all datasets, we performed 631 statistical comparisons for the expression levels of the 11 canonical brain shuttle targets (TFRC, SLC3A2, IGF1R, TMEM30A, ALPL, SLC2A1, INSR, LDLR, LRP1, LRP8, CA4). In the main figures, we highlight five representative targets, TFRC, SLC3A2, IGF1R, TMEM30A, and ALPL, that together exemplify distinct endogenous BBB transport pathway classes, with results for the remaining six targets provided in the Supplementary Figures. Of the 631 comparisons, 612 (97%) showed no significant difference (Supplementary Table 1). The remaining 19 comparisons (3%) that reached statistical significance, together with the non significant comparisons, are described in the subsections below.

### Brain shuttle targets show no regional or Alzheimer’s disease-related differences

We next focused on AD to examine regional and disease related variations in the expression of specific transport target proteins and to determine whether individual receptor levels differed between conditions. BMEC snRNA-seq data from two published studies ^35, 36^ were analyzed at the individual patient level (Fig. 2). The data included samples from the entorhinal cortex (EnC), inferior temporal gyrus (ITG), prefrontal cortex (PFC), and visual cortex (V1/V2), and from the superior frontal cortex (SFC) and hippocampus (HPC), respectively, in both AD and control subjects (Fig. 2a). For every patient within each study, transcript expression was calculated for all cell surface membrane protein-encoding genes, and data for the eleven brain shuttle target genes were selected for detailed analysis, with data for five representative targets presented here (Figure 2). All but two comparisons detected no significant differences in expression between regions or disease states. The only exception was seen in SLC3A2 expression, which was significantly higher in AD V1 samples compared to AD PFC and ITG samples. Analysis of the six additional canonical brain shuttle targets revealed some regional differences within the AD group (Supplementary Fig. 1). LRP1 showed the most pronounced regional variation, with significantly higher expression in AD PFC compared to all four other AD regions. Both LDRL and INSR exhibited significantly higher expression in AD V2 compared to AD EnC and PFC, respectively. INSR also showed a significant difference in the control group, with higher expression in Control V1 compared to Control EnC. No significant regional or disease-related differences were observed for these genes in the SPC or HPC regions. Notably, including SLC3A2 from the main figure, significant regional differences were predominantly observed within the AD group rather than the control group. Together with the main figure results, SLC2A1, LRP8, and CA4 showed no significant differences in any comparison, and overall, transcript expression of canonical brain shuttle targets in BMECs remains largely consistent across brain regions and between AD and control samples, as only 11 of the 319 (3.4%) post-hoc pairwise comparisons performed in this analysis exhibited statistically significant differences (Supplementary Table 1).

**Fig. 2:**
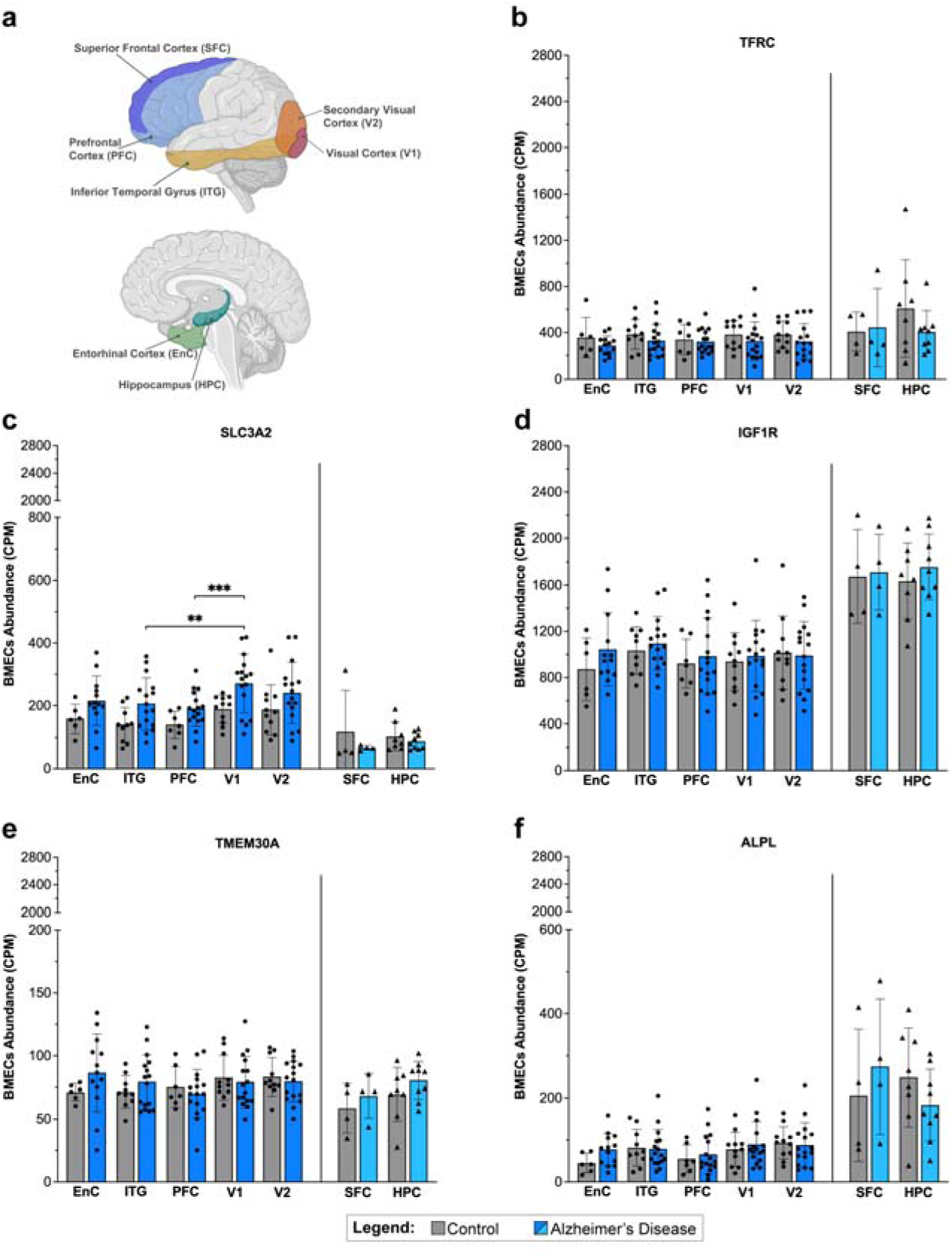
Canonical brain shuttle target genes show very few significant expression differences between brain regions or disease states in Alzheimer’s and control samples. (**a**) Schematic of the brain highlighting regions analyzed: entorhinal cortex (EnC), inferior temporal gyrus (ITG), prefrontal cortex (PFC), visual cortex (V1/V2), superior frontal cortex (SFC), and hippocampus (HPC). (**b-f**) Patient-level expression of five representative BMEC surface BBB transport genes measured by snRNA-seq in Alzheimer’s disease (AD) and control subjects, from two independent studies. Data from ^36^ (circles) include EnC (n = 19, 6:13 Control:AD), ITG (n = 26, 10:16 Control:AD), PFC (n = 23, 7:16 Control:AD), and V1/V2 (n = 27, 11:16 Control:AD); data from ^35^ (triangles) include SFC (n = 8, 4:4 Control:AD) and HPC (n = 17, 8:9 Control:AD). Expression (CPM) plotted by patient, region, and disease state. Linear mixed-effects models (Disease × Region, with PatientID as random intercept) were performed within each dataset, with post-hoc pairwise comparisons via estimated marginal means with Tukey adjustment; graphs show mean ± SD, **p≤0.01, ***p≤0.001, Benjamini-Hochberg (BH)-corrected for all genes tested.

### Brain shuttle target expression shows very few differences related to additional regions or diseases including PD, ALS and HD

Having established that brain shuttle target expression is largely similar between regions and diseases states in AD and control subjects, we next applied the same patient level BMEC snRNA seq analysis to other neurodegenerative diseases and their characteristic affected regions. Specifically, we analyzed three additional datasets: midbrain in PD ^38^, motor cortex in c9ALS and sALS ^40^, and caudate/putamen in HD ^39^, each with matched controls (Fig. 3). For each patient, transcript expression was first calculated for all cell surface membrane protein-encoding genes, and data for the eleven canonical brain shuttle target genes were extracted for comparison.

**Fig. 3:**
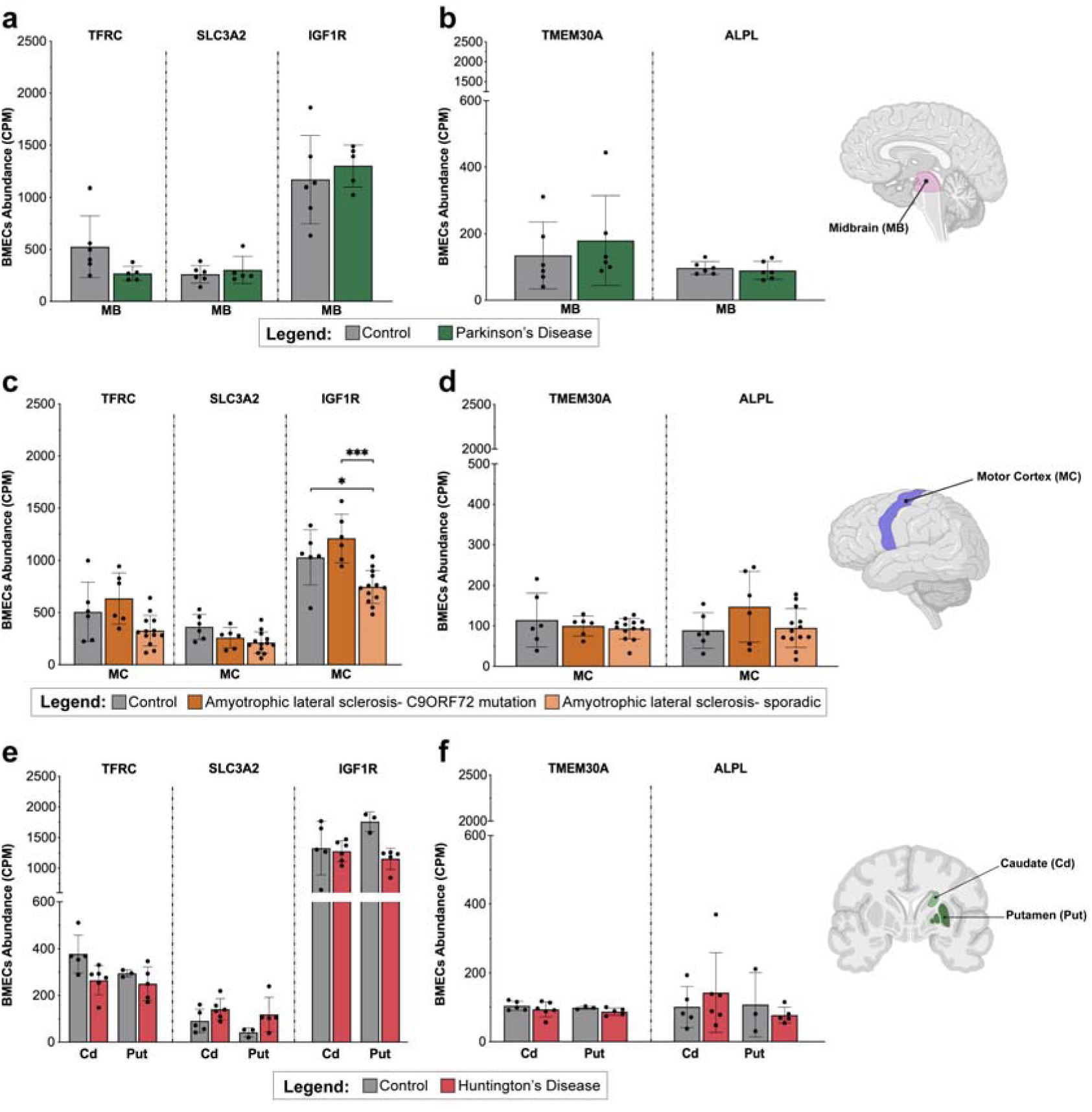
Brain shuttle target transcript expression is conserved between brain regions and between neurodegenerative disease and control subjects. (**a–f**) Expression of five brain shuttle targets in brain microvascular endothelial cells (BMECs), quantified by snRNA-seq in three independent datasets (CPM, counts per million). Genes shown include TFRC, SLC3A2/CD98hc, and IGF1R (**a, c, e**) as well as TMEM30A and ALPL (**b, d,f**). Brain regions analyzed are indicated by schematic insets in each panel. (**a, b**) Midbrain (MB) from control (grey, n = 6) and Parkinson’s disease (PD; green, n = 5) patients. (**c, d**) Motor cortex (MC) from control (grey, n = 6), C9ORF72-associated ALS (c9ALS; dark orange, n = 6), and sporadic ALS (sALS; light orange, n = 13) patients. (**e, f**) Caudate (Cd) and putamen (Pt) from control (Cd: n = 4, Pt: n = 3) and Huntington’s disease (HD; red; Cd: n = 6, Pt: n = 5) patients. Expression plotted per patient by region and disease status. Welch’s t-test was used for Smajic et al. (Control vs. PD: Panels a and b); one-way ANOVA with Tukey’s HSD post-hoc test was used for Pineda et al. (Control vs. c9ALS vs. sALS: Panels c and d); and a linear mixed-effects model (Disease × Region, with PatientID as random intercept) with post-hoc pairwise comparisons via estimated marginal means with Tukey adjustment was used for Garcia et al. (Control vs. HD: Panels e and f). All p-values were BH-corrected over all tested genes; graphs show mean ± SD, *p≤0.05, ***p≤0.001.

In midbrain samples from PD and control subjects, gene expression of all five brain shuttle targets was unchanged between groups (Fig. 3a, b). In motor cortex samples from c9ALS, sALS, and control subjects, only IGF1R was significantly lower in sALS compared to both control and c9ALS. No significant differences were observed for TFRC, SLC3A2, TMEM30A, or ALPL (Fig. 3c, d). In striatum samples from HD and control subjects, all of the assessed genes showed no significant changes (Fig. 3e, f). Analysis of the additional canonical brain shuttle targets showed no significant differences between any groups (Supplementary Fig. 2), except for LRP1, which was significantly increased in motor cortex samples from sALS compared to both control and c9ALS. Overall, these results demonstrate that BMEC transcription of brain shuttle targets remains highly stable between diverse brain regions and largely preserved between neurodegenerative disease states.

### Brain shuttle target expression is stable between males and females and between control and AD subjects

We next asked whether brain shuttle target expression varies between males and females in AD, and whether such differences are detectable at both the transcript and protein level. To assess the variation in brain shuttle target transcript and protein expression in male versus female and AD versus control groups, we analyzed multiple transcriptomic and proteomic datasets. scRNA-seq, snRNA-seq, and proteomic data for the eleven canonical brain shuttle target genes were examined in BMECs and isolated human brain microvessels (BMVs) (Fig. 4, Supplementary Fig. 3). All samples in the three independent scRNA-seq studies ^41-43^ were from healthy subjects; these datasets were therefore used exclusively to compare expression between males and females (Fig. 4a, Supplementary Fig. 3a). No significant differences in brain shuttle target expression were detected between sexes in these datasets.

**Fig. 4:**
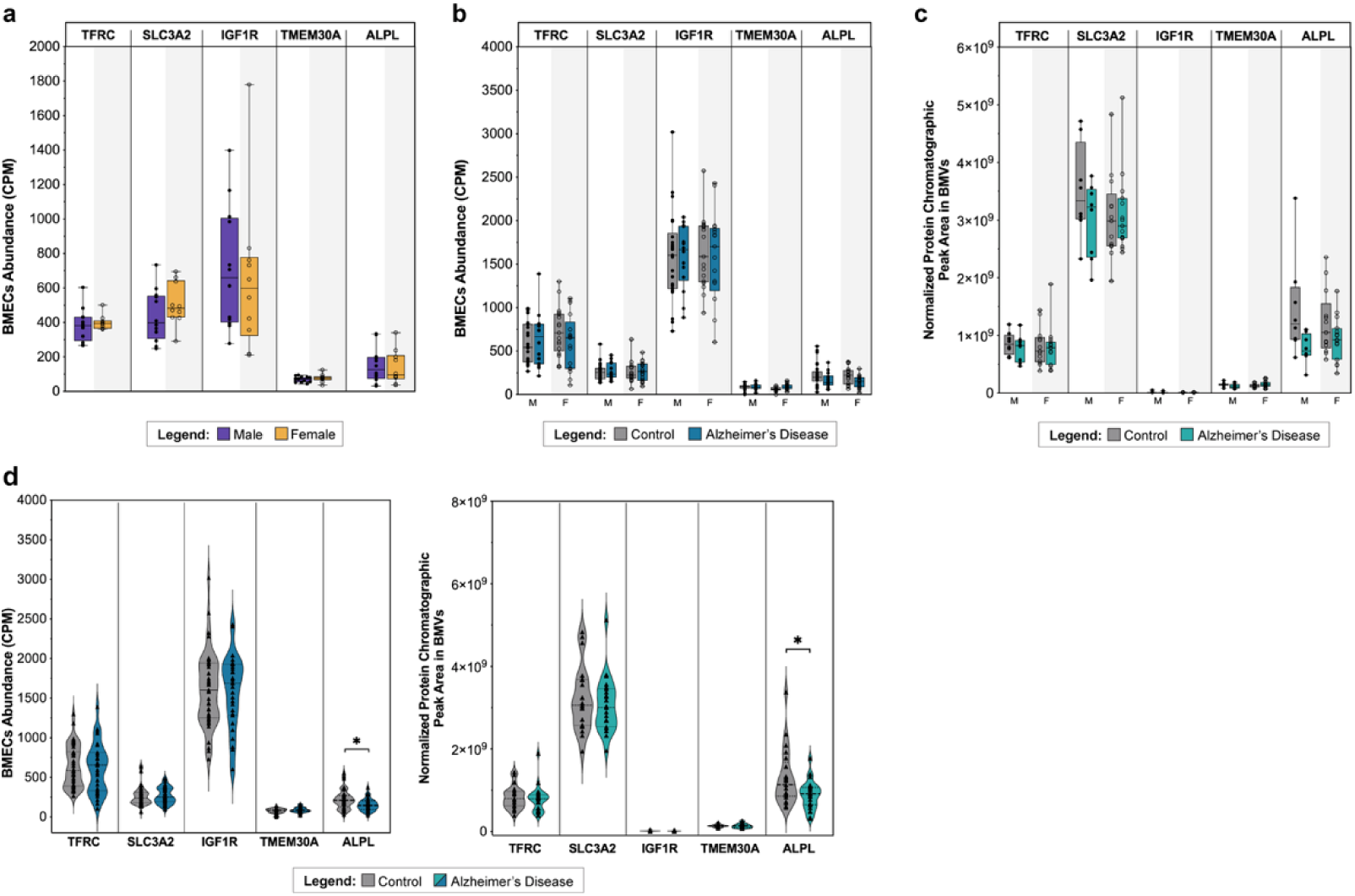
Brain shuttle target transcript and protein expression are conserved between males and females and between control and Alzheimer’s disease subjects. (**a-c**) Expression of five brain shuttle genes in brain microvascular endothelial cells (BMECs) or brain microvessels (BMVs), stratified by sex (males: filled black circles; females: open circles) and Alzheimer’s disease (AD) status (control: grey bars, AD: blue/teal bars). (**a**) scRNA-seq data from ^41-43^ for healthy controls (purple bars: males, n = 12; yellow bars: females, n = 10). (**b**) snRNA-seq data from ^37^ for control (grey) and AD (blue) subjects: control males (n = 26), AD males (n = 14), control females (n = 17), AD females (n = 17). Transcript levels are shown as counts per million (CPM). (**c**) Proteomic data from ^44^ showing normalized protein abundance in BMVs from the superior frontal cortex in control (grey) and AD (teal) subjects: control males (n = 8), AD males (n = 8), control females (n = 13), AD females (n = 15). (**d**) For both snRNA-seq ^37^ and proteomic datasets ^44^, additional analyses of differences between control and AD groups were performed with males and females combined. Graphs display individual values and mean ± SD. A linear mixed-effects model (Expression ∼ Sex + (1|Study)) was used for the scRNA-seq data (panel a); two-way ANOVA (Disease × Sex) with Tukey’s HSD post-hoc test was used for Sun et al. and Erickson et al. sex analyses (panels b and c, respectively); and Welch’s t-test was used for Sun et al. and Erickson et al. disease-only comparisons (panel d). All p-values were BH-corrected over all tested genes; graphs show mean ± SD, *p≤0.05.

For disease versus healthy control comparisons, snRNA-seq ^37^ and proteomic ^44^ data were used, allowing us to compare sexes in AD and control groups (Fig. 4b, c). These datasets were analyzed both with sexes separated (Fig. 4b, c, Supplementary Fig. 3b, c) and with sexes combined (Fig. 4d, Supplementary Fig. 3d). Of the eleven transport targets investigated in this study, two proteins, LRP8 and LDLR, were not detected in the proteomic dataset. In the analyses considering both sex and disease status (Fig. 4b, c, Supplementary Fig. 3b, c), none of the genes showed significant post-hoc pairwise differences in expression between any of the four group comparisons (Control Male vs. Control Female, AD Male vs. AD Female, Control Male vs. AD Male, Control Female vs. AD Female), at either the transcript or protein level. In the comparison of AD versus control with sexes combined (Fig. 4d, Supplementary Fig. 3d), ALPL exhibited a significant reduction in AD versus control subjects at both transcript and protein levels, while the proteomic analysis alone showed a significant reduction in LRP1 expression in AD. Thus, reduced ALPL protein and transcript abundance in AD is the only consistent, significant change among the analyzed endothelial surface proteins when comparing AD and control samples.

To further test these findings, we analyzed one additional proteomic dataset was analyzed. In this dataset ^45^, only ALPL showed significantly lower expression in AD compared to control (p < 0.05, |log FC| ≥ 1), while expression of other brain shuttle targets did not differ significantly (Supplementary Table 2). These consistent results between multiple transcriptomic and proteomic datasets provide striking support for the conservation of transport target protein expression between males and females and healthy versus AD subjects, with ALPL being the sole exception.

### Brain shuttle target expression remains stable with age

We then investigated whether expression of brain shuttle targets varies with age. To determine whether age is correlated with brain shuttle target transcript or protein expression, we analyzed multiple transcriptomic and proteomic datasets (Fig. 5). Expression of the eleven canonical targets was assessed in BMECs or BMVs using scRNA-seq, snRNA-seq, and proteomic data. In healthy control scRNA-seq datasets ^41-43^, individuals were stratified into three age categories: 15-29, 30-45, and over 45 years; these age ranges were selected to obtain balanced group sizes so that statistical comparisons would be adequately powered, with the trade off that age bins are not identical between all datasets. Analyses of transcript levels between these groups revealed no significant age-related differences for any genes (Fig. 5a). For snRNA-seq datasets ^35, 36^, groups were defined as 55-84 years and 85+ years to divide middle-aged and older individuals into similarly sized groups, again prioritizing sufficient and balanced sample sizes for statistical testing and relevance for AD comparisons (Fig. 5b). For the proteomic data ^44^, we utilized two groups 58-89 years and 90+ years, reflecting the age distribution in this cohort (Fig. 5c). In both the snRNA-seq and proteomic analyses, no significant age-associated changes were detected for the five brain shuttle target proteins in control or AD subjects.

**Fig. 5:**
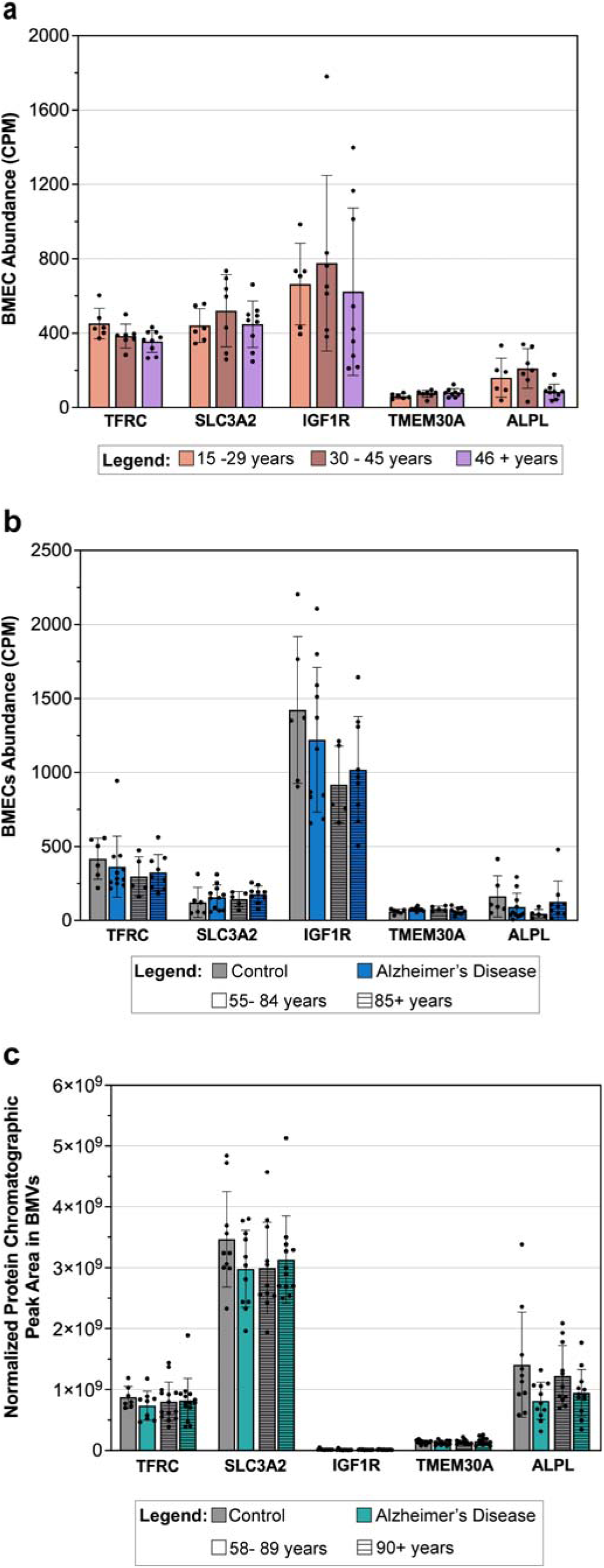
Brain shuttle gene expression transcript and protein levels are conserved in different age groups. (**a–c**) Expression of five brain shuttle transport genes in brain microvascular endothelial cells (BMECs) or brain microvessels (BMVs), stratified by disease status and age, using sc/snRNA-seq or proteomics. (**a**) scRNA-seq data from healthy control datasets ^41-43^, showing BMEC transcript levels (CPM) in individuals aged 15–29 (n = 6), 30–45 (n = 7), and >45 (n = 9). (**b**) snRNA-seq data ^35, 36^, combining prefrontal (PFC) and superior frontal cortex (SFC) samples. Control (grey) and AD (blue), solid bars indicate 55–84 years and horizontally striped bars indicate ≥85 years (control: n = 6/5; AD: n = 11/9, respectively). (**c**) Proteomic data ^44^, showing normalized protein abundance in BMVs from SFC of Control (grey) and AD (teal). Solid bars indicate 58–89 years and horizontally striped bars indicate ≥90 years (control: n = 10/11; AD: n = 11/12, respectively). A linear mixed-effects model (Expression ∼ AgeGroup + (1|Study)) with post-hoc pairwise comparisons via estimated marginal means with Tukey adjustment was used for the scRNA-seq data (panel a); a linear mixed-effects model (Disease × AgeGroup + (1|Study)) with Tukey-adjusted post-hoc comparisons was used for Yang/Bryant et al. (panel b); and two-way ANOVA (Disease × AgeGroup) with Tukey’s HSD post-hoc test was used for Erickson et al. (panel c). All p-values were BH-corrected; graphs show mean ± SD; all statistical comparisons were non-significant.

The six additional brain shuttle target genes (SLC2A1, LRP1, LRP8, LDLR, INSR, CA4) showed no significant differences by age or disease status, except for a decrease in LRP1 protein expression in AD compared to control individuals in the younger cohort in the proteomic dataset. (Supplementary Fig. 4). Together, these results demonstrate that expression of the canonical brain shuttle targets remains remarkably stable regardless of age and AD status at both the transcript and protein levels.

### Brain shuttle target transcript and protein expression varies significantly between individuals

Throughout the preceding analyses, brain shuttle targets remarkably similar expression between groups defined by brain region, neurodegenerative disease status, sex, or age. In striking contrast, we observed high variability between individual subjects within every group analyzed. To formally quantify this, we performed variance decomposition for every analysis in Figures 2-5 (full results in Supplementary Data File 2). Between all datasets and analyses, the residual, representing variance not attributable to the fixed effects tested, including inter-individual biological variability, measurement noise, and, in mixed-effects models, variance associated with the random effects, was overwhelmingly the dominant component, accounting for a median of 94.8% of total variance in gene expression. In over 91% (125/137) of gene-by-analysis combinations, the residual explained ≥80% of the variance. By comparison, the factors tested (brain region, disease status, sex, age, and their interactions) collectively explained only a small fraction of expression variance in most cases (Table 1, Supplementary Data File 2).

**Table 1:** Inter-individual variation in expression of brain shuttle target genes in transcriptomic and proteomic datasets. (**a–c**) Expression of five brain shuttle target genes was assessed in three datasets: (**a**) scRNA-seq data from control individuals (n = 22) ^41-43^, (**b**) snRNA-seq data from control (n = 43) and AD (n = 31) individuals ^37^, and (**c**) proteomic data from control (n = 21) and AD (n = 23) individuals ^44^. Within each group, patients were stratified as high or low expressors based on the group median. The table reports, for each gene, the expression range, foldDdifference between highest and lowest expressors, mean expression in high vs. low expressors, and variance explained by residual for each analysis conducted for each dataset.

| Symbol | Expression Range [CPM] | Fold Difference (Highest/Lowest Ind) | Mean high Expressors [CPM] | Mean Low Expressors [CPM] | % Variance explained by Residual (Fig 4a) | % Variance explained by Residual (Fig 5a) |
| --- | --- | --- | --- | --- | --- | --- |
| *TFRC* | 266-432 | 1.6 | 405 | 304 | 97.53 | 71.7 |
| *SLC3A2* | 248-661 | 2.7 | 544 | 347 | 93.73 | 86.44 |
| *IGF1R* | 211-1399 | 6.6 | 1031 | 266 | 99.42 | 99.91 |
| *TMEM30A* | 53-124 | 2.4 | 98 | 63 | 97.81 | 82.94 |
| *ALPL* | 36-177 | 4.9 | 115 | 57 | 99.43 | 76.55 |

**b**
| Symbol | Disease State | Expression Range [CPM] | Fold Difference (Highest/Lowest Ind) | Mean high Expressors [CPM] | Mean Low Expressors [CPM] | % Variance explained by Residual (Fig 4b) | % Variance explained by Residual (Fig 4d – Left Panel) |
| --- | --- | --- | --- | --- | --- | --- | --- |
| *TFRC* | Healthy | 267-1300 | 4.9 | 867 | 418 | 95.67 | 99.78 |
|  | AD | 109-1389 | 12.8 | 875 | 352 |  |  |
| *SLC3A2* | Healthy | 65-636 | 9.8 | 352 | 178 | 99.46 | 99.96 |
|  | AD | 95-486 | 5.1 | 358 | 181 |  |  |
| *IGF1R* | Healthy | 729-3019 | 4.1 | 1981 | 1234 | 99.68 | 99.86 |
|  | AD | 603-2432 | 4.0 | 1951 | 1185 |  |  |
| *TMEM30A* | Healthy | 0-145 | Inf | 101 | 49 | 90.63 | 95.72 |
|  | AD | 21-161 | 7.8 | 116 | 63 |  |  |
| *ALPL* | Healthy | 29-556 | 19.4 | 315 | 132 | 89.99 | 90.31 |
|  | AD | 22-370 | 17.2 | 216 | 93 |  |  |

**c**
| Symbol | Disease State | Expression Range [Normalized Peak Area E+05] | Fold Difference (Highest/Lowest Ind) [Normalized Peak Area E+05] | Mean High Expressors [Normalized Peak Area E+05] | Mean Low Expressors [Normalized Peak Area E+05] | % Variance explained by Residual (Fig 4c) | % Variance explained by Residual (Fig 4d – Right panel) | % Variance explained by Residual (Fig 5c) |
| --- | --- | --- | --- | --- | --- | --- | --- | --- |
| *TFRC* | Healthy | 3879 - 14396 | 3.7 | 10532 | 6042 | 98 | 99.19 | 90.08 |
|  | AD | 3903 - 18913 | 4.9 | 10007 | 5742 |  |  |  |
| *SLC3A2* | Healthy | 19447 - 48363 | 2.5 | 38421 | 26142 | 93.8 | 98.96 | 92.53 |
|  | AD | 19629 - 51251 | 2.6 | 35795 | 25460 |  |  |  |
| *IGF1R* | Healthy | 71 - 521 | 7.4 | 181 | 100 | 92.05 | 98.23 | 91.66 |
|  | AD | 54 - 421 | 7.9 | 162 | 89 |  |  |  |
| *TMEM30A* | Healthy | 810 - 2201 | 2.7 | 1640 | 1103 | 92.2 | 99.84 | 94.82 |
|  | AD | 726 - 2583 | 3.6 | 1826 | 952 |  |  |  |
| *ALPL* | Healthy | 5828 - 33798 | 5.8 | 18248 | 8149 | 82.07 | 84.94 | 79.11 |
|  | AD | 3141 - 17707 | 5.6 | 11564 | 6090 |  |  |  |

The magnitude of this inter-individual variation is further illustrated by descriptive measures of expression spread within groups (Table 1, Supplementary Table 3). Within each dataset, individuals were stratified as high or low expressors based on the group median for each gene. Fold differences between the highest and lowest expressing individuals within the same group routinely ranged from 2- to 10-fold, and in some cases exceeded an order of magnitude, far exceeding any differences attributable to brain region, disease, sex, or age. This pattern was consistent across all 11 brain shuttle targets, across both transcriptomic and proteomic datasets, and across healthy control and disease cohorts.

To determine whether this inter-individual variation might be an artifact of post-mortem interval (PMI), defined as the delay between time of death and harvesting of organs, we performed Spearman correlation analysis between PMI and brain shuttle target expression levels for subjects in the Erickson proteomic dataset^44^. No significant correlation between PMI and expression level was detected in any dataset for any of the genes examined (Supplementary Fig. 5), indicating that PMI does not explain the substantial inter individual variation in brain shuttle target expression.

## DISCUSSION

Our findings directly address a critical but previously unexamined assumption underlying the clinical development of brain shuttle therapeutics: that the endothelial receptor targets on which these strategies depend are sufficiently and stably expressed between individuals and patient groups. We anticipated that expression of brain shuttle targets would vary substantially with disease state, brain region, age, and sex, while remaining relatively consistent between individuals within the same group. Our data revealed the opposite. By integrating transcriptomic and proteomic data from large human cohorts, we find that brain shuttle target expression is remarkably stable across disease states, brain regions, sexes, and ages. Of 631 statistical comparisons, only 19 (3%) revealed significant differences. In contrast, inter-individual variation was significant in every group and for every target examined. Together, these results strongly suggest that individual biology, rather than anatomy, disease, or demographics, is the dominant and most clinically relevant source of variation in brain shuttle target expression.

Most previous studies of regional protein expression in neurodegenerative diseases ^46-58^ have analyzed transcriptomic or proteomic data to characterize molecular changes in bulk brain tissue and parenchymal cell populations between healthy and neurodegenerative conditions, such as AD, PD, ALS, and HD, often highlighting region- or disease-specific alterations ^46-51^. These studies, primarily reflecting signals from neurons and glia, have shown significant shifts in gene and protein expression between regions and disease states, including changes in stress responses, immune pathways, and sex-specific gene expression, in humans and mice ^46-58^. In contrast, when we analyzed transcriptomic datasets of healthy controls and AD, PD, ALS and HD patients to specifically examine the BMEC expression of brain shuttle targets in different brain regions, we found that brain shuttle target expression is largely stable between brain regions, with only a small number of region-specific differences observed. Within each dataset, comparisons of brain regions in healthy controls or diseased subjects showed consistent transcript and/or protein expression levels for every target. The discrepancy between our findings and those of past studies likely occurs because we focused on the expression of specific brain shuttle targets in BMECs and BMVs rather than in bulk tissue samples, allowing us to analyze endothelial-specific expression that is diluted and obscured in whole-tissue expression analyses dominated by neuronal and glial transcripts or proteins. One mouse study, which used scRNA-Seq to analyze regional transcript level variation in BBB genes, reported that wild-type male mice showed increased endothelial TFRC expression in the hippocampus and striatum, suggesting the potential for region-specific TFRC-mediated drug delivery ^59^. In contrast, our analysis of TFRC regional transcript expression between multiple large human datasets encompassing control, AD, PD, ALS, and HD cohorts found minimal (11/319 post-hoc comparisons) significant regional variations within any dataset. This difference in results may reflect species differences in regional expression. Such species differences are a well-recognized issue in preclinical-to-clinical translation ^60-62^, and our findings underscore why analyses of large multi-omics human datasets are critical for guiding the development of any brain shuttle-linked therapeutics, for which delivery efficiency may be significantly altered by variation in shuttle target expression.

Importantly, we also found no variation in BMEC or BMV expression of brain shuttle targets between healthy controls and diseased patients in the four neurodegenerative disease datasets analyzed: AD, PD, ALS and HD, except for decreased ALPL transcript and protein, and LRP1 protein expression in AD compared to control. The question of whether brain shuttle target expression is substantially maintained in diseased patients, in whom the BBB is known to undergo disease-related changes in structure and function ^33, 34^, is critical when drug efficacy depends on brain shuttles to mediate enhanced drug uptake in these patients. Previous studies of these target proteins in healthy versus diseased subjects have focused on elucidation of altered functions such as disrupted TfR-mediated iron regulation ^63, 64^, impaired IGF1R/INSR signaling ^65, 66^, perturbation of TMEM30A interaction with APP β-CTF and of endosomal trafficking ^67^, increased ALPL activity in both hippocampus and plasma of AD patients compared with controls ^68^, and broader changes in metabolic or transporter pathways in diverse neurodegenerative diseases ^69-71^. Our study uses multi-omics and the resulting statistical power to assess expression levels of brain shuttle targets in BMECs or BMVs in healthy vs diseased populations. Our findings establish that in the four major neurodegenerative diseases analyzed, target expression levels do not differ significantly between healthy and diseased subjects. For clinical development of drug-shuttle fusions, this finding provides important reassurance that target receptor abundance at the BBB is preserved in subjects with neurodegenerative diseases.

In our analysis of expression by sex, none of the eleven brain shuttle targets showed significant differences between males and females in any dataset or disease group examined. While sex differences in BBB transporter expression on LRP1 and SLC2A1 have been reported in rodent models ^72^, a recent in vitro study of primary mouse brain endothelial cells found no significant sex differences in TfR, INSR and SLC2A1 BBB transporter expression ^73^. Systematic characterization of sex-dependent variation in brain shuttle target expression in human brain tissue has been limited ^74^, and our findings in large human cohorts do not support sex as a meaningful source of variation in this context. It should be noted that our study was not designed to detect acute temporal variation in expression, such as potential hormonally mediated changes between different phases of the menstrual cycle.

Our analysis of expression variation by age showed no significant differences for a large majority of comparisons between all eleven targets. Prior studies have reported age-related changes in several brain shuttle targets in animal models and human tissues, including declining LRP1 and GLUT1 at the aging brain endothelium and contradictory findings for TFRC ^25, 72, 75-77^. The only significant difference in our dataset was a small but significant reduction in LRP1 protein expression between healthy controls and AD patients within the 58-89 age group in the proteomic dataset only, not replicated in transcriptomic datasets or in the 90+ age group. The overall stability of expression between age groups for all eleven brain shuttle targets confirms that aging is not associated with substantially decreased expression of most, and likely all, human brain shuttle targets tested, an important reassurance for clinical indications targeting older patient populations.

The overall stability of brain shuttle target expression across brain regions, disease states, sex, and age is itself a meaningful finding for the field. It indicates that brain shuttle strategies are unlikely to require anatomical, disease-specific, or demographic tailoring of their target selection, simplifying one dimension of clinical trial design and broadening the potential applicability of a given shuttle beyond limited patient populations and disease stages.

In striking contrast to this uniformity, we found highly significant inter-individual variation in brain shuttle target expression in all groups analyzed. Brain shuttle target abundance varied between the lowest- and highest-expressing individuals more than an order of magnitude. Critically, this inter-individual variability, as measured by both variance decomposition (Supplementary Data File 2) and descriptive expression spread (Table 1, Supplementary Table 3) was observed for all eleven brain shuttle targets, in both healthy controls and AD patients, and was consistent between all three data modalities, scRNA-seq, snRNA-seq, and proteomic datasets, making it the single most robust and reproducible finding of this study. To our knowledge, this pronounced inter-individual variation, though a well-recognized feature of biological systems, has not previously been reported in the context of brain shuttle target expression or discussed as an important factor in the design and interpretation of clinical trials for new shuttle-fused large molecule therapeutics ^78, 79^. The clinical implications are direct: patients with the lowest receptor expression level might achieve substantially less brain exposure than those with the highest target expression at identical administered doses, a source of pharmacokinetic heterogeneity that is invisible to any trial design that does not measure and account for brain transport target expression. While previous research has focused on improving drug delivery and stratifying patients with disease-related biomarkers ^80, 81^, individual differences in brain shuttle target expression may critically influence the uptake of, and therapeutic response to, drugs using brain shuttles to enhance brain uptake. This has particular relevance for interpreting the variable clinical responses observed between patients enrolled in the TfR-targeted brain shuttle program trontinemab ^82^, and the heterogeneous CNS exposure outcomes reported in clinical trials of TfR-fused enzyme replacement therapies ^17, 18^. Thus, there is an urgent need to identify and validate clinical biomarkers that quantify brain shuttle target abundance at the individual level, analogous to HER2 ^83, 84^ or EGFR ^84^ expression testing in oncology, to enable patient stratification and management. Given the absence of routine biopsies of brain microvasculature, the development of biomarkers presents a significant challenge, possibly relying on PET or other types of imaging. However, just as improved imagine of brain pathology and measurement of disease biomarkers have improved patient selection in Alzheimer’s trials, the availability of biomarkers for brain shuttle target expression could greatly improve patient selection and dosing for drugs fused to brain shuttles in diverse patient groups.

Notably, expression differences for LRP8 and LDLR were consistently detected at the RNA level but not at the protein level. Such RNA-protein discordance is well recognized and arises from both technical factors, including the lower sensitivity of proteomic methods for low-abundance or hydrophobic membrane proteins along with batch effects from tissue processing ^85, 86^, and biological factors such as differences in translation efficiency, protein stability, and post-translational regulation ^86, 87^. These findings reinforce the value of the multi-omics approach used here: convergence between modalities, as observed for the nine targets detected at both RNA and protein levels, provides the strongest evidence for the conclusions of stable expression drawn in this study.

As with any large-scale multi-cohort study, several considerations should be noted when interpreting our findings. Age categories were determined by sample distributions and the need for adequate group sizes, resulting in categories that differ between datasets and omics types; while this ensured appropriate statistical power within each analysis, direct cross-dataset comparisons of age effects are not possible. In each case, our findings allow the generation of hypotheses that should be further tested in larger cohorts. Although we used the largest available human datasets, testing of additional brain regions and disease states will be possible as additional datasets emerge. Our proteomic analyses relied heavily on BMV preparations, which include not only BMECs but also variable degrees of contaminating adherent astrocyte endfeet, pericytes, and microglia. While this precludes the possibility of pure cell-type-specific quantification, BMV proteomics nonetheless provides a substantially more accurate estimate of BMEC protein abundance than bulk brain tissue and constitutes a powerful complement to our transcriptomic data. Finally, most of the dataset analyzed were obtained from post-mortem tissue; while the fact that we found no correlation between PMI and target protein abundance strongly suggests that PMI is not a key determinant of measured abundance levels, we cannot fully exclude potential influences of rapid post-mortem changes on protein expression.

Future work should extend these analyses to additional populations and diseases, including the use of spatial transcriptomics along with single-cell and single molecule proteomics for greater anatomical and cellular resolution. Directly linking transporter abundance to in vivo CNS drug exposure or therapeutic outcomes will be important to establish whether the inter-individual variability in target expression we observe translates into clinically meaningful differences in brain drug exposure. Pre-clinical studies in mice engineered to express varying levels of brain shuttle target proteins could provide one model to quantify the impact of transporter expression on brain pharmacokinetics, target engagement, and efficacy. Most urgently, the development of companion biomarkers such as in vivo imaging assays to quantify brain shuttle target protein expression levels in human subjects would enable improved patient selection, stratification and management of response to treatment with drugs linked to brain shuttles.

## MATERIALS AND METHODS

### Surface protein list generation

Surface protein–coding genes were defined as those encoding either TM proteins or GPI-APs. TM proteins were identified through an integrated approach combining protein domain annotation with selected related UniProt and Gene Ontology (GO) keywords. GO terms included: “plasma membrane”, “integral component of plasma membrane”, “cell surface”, “intrinsic component of plasma membrane”, “apical plasma membrane”, “apicolateral plasma membrane”, “basal plasma membrane”, and “basolateral plasma membrane.” UniProt terms included: “cell membrane”, “cell surface”, “apical cell membrane”, “apicolateral cell membrane”, “basal cell membrane”, and “basolateral cell membrane.” Genes were classified as encoding surface proteins if their corresponding protein was annotated with a transmembrane domain and at least one relevant keyword match, or if the protein contained at least one extracellular topological domain. This process yielded a total of 3,527 TM proteins.

GPI-anchored proteins were defined based on the presence of the “GPI” keyword in UniProt, supplemented by cross-referencing with the most frequent GO annotations found among GPI-APs (notably “plasma membrane” and “side of membrane”). Proteins were excluded if they were not conventional GPI-APs or if GPI-anchor status could not be confirmed using Protter (https://wlab.ethz.ch/protter/start/). This filtering resulted in the identification of 137 GPI-APs. By combining these strategies, we generated a comprehensive list of genes encoding surface-exposed TM proteins and GPI-APs for downstream analysis (Supplementary Data File 1).

### sc/snRNAseq data alignment and analysis

Raw sequencing data (BAM/FASTQ files) for the following publications were obtained from GEO and dbGaP repositories: for scRNA-seq (healthy brain), Winkler et al. 2022 ^41^ (dbGaP phs002624.v2.p1), Xie et al. 2024 ^42^(GSE242044), and Wälchli et al. 2024 ^43^(GSE256493); and for snRNA-seq (healthy and diseased brain), Yang et al. 2022 ^35^(GSE163577), Bryant et al. 2023 ^36^(PRJNA916657), Garcia et al. 2022 ^39^(GSE173731), Smajić et al. 2022 ^38^(GSE157783), and Pineda et al. 2022 ^40^(PRJNA1073234). For Sun et al. 2023^37^, pre-aligned data provided by the authors (https://compbio.mit.edu/scADbbb/) were reanalyzed; therefore, data from this study were not included in the correlation analysis in Fig. 1. Patient demographics, sample numbers, and brain region details for each dataset are provided in Supplementary Data File 3.

In instances where only BAM files were available, the raw BAMs were converted to FASTQ format using the bamtofastq utility (version 1.3.2). The resulting FASTQ files, along with those already provided in FASTQ format, were processed using Cell Ranger (version 7.0.0; 10x Genomics) to perform read alignment and generation of the gene expression count matrix. For all datasets, FASTQ files were aligned to the GRCh38 human reference genome (refdata-gex-GRCh38-2020-A; 10x Genomics) with intronic reads included by default to accommodate the nuclear RNA content of snRNA-seq.

All analyses were performed independently for data from each publication. Within each study, raw count matrices from all samples were imported into R (Seurat v4.3) and merged into a single Seurat object. Quality control thresholds, including minimum gene and molecule counts per cell and maximum allowable mitochondrial read proportion, followed the standards set in the original publications. Only cells passing these criteria were retained. Notably, gene filtering by minimum cell detection (“min.cells”) was not applied, so all 36,601 genes were included in downstream analyses.

After normalization and scaling, highly variable genes were identified, and principal component analysis (PCA) was run using the top 2,000 variable genes. The number of significant principal components to retain was determined using the Elbow plot heuristic, resulting in the use of 30 PCs for downstream steps. UMAP visualization was then performed using these 30 PCs.

Cell clustering was performed in Seurat using the FindClusters function (resolution 1.5), deliberately producing granular clusters to facilitate clear distinction of major endothelial subtypes (arterioles, capillaries, venules, veins, and lymphatics) as well as non-endothelial cell populations (astrocytes, neurons, oligodendrocytes, pericytes, and microglia/macrophages) present in the dataset. BMEC clusters were then specifically identified by their selective expression of established BMEC/capillary markers (e.g., SLC12A1, NPIPB5, MFSD2A, SLC7A5, SLC16A1, INSR, IVNS1ABP), and these clusters were further refined to ensure exclusion of other endothelial and contaminating cell types.

This over-clustering and subsequent targeted identification approach enables more accurate delineation of BMECs compared to direct low-resolution clustering ^88^. For each study, the finalized BMEC clusters were used to calculate counts per million (CPM) values for each gene. Aggregate CPM values were then calculated for all patients within a given disease status or brain region group; these aggregates were used to perform Spearman correlation analysis of surface protein expression profiles and to assess similarity between different diseases and brain regions. Additionally, patient level CPM values were generated for each gene and used in subsequent analyses examining differences in key brain shuttle target expression by sex, age, neurodegenerative disease status, and brain region.

### Proteomic data preprocessing and analysis

Proteomic data from two independent studies^44, 45^ underwent dataset-specific preprocessing. For ^44^, normalized protein chromatographic peak areas were used as the starting values. Zeros were first replaced with the minimum nonzero value for each sample. Data were then median-normalized so that the median abundance for each sample was scaled to match a global target median calculated between all samples. This processed dataset was then used for all downstream analyses of brain shuttle target expression. For ^45^, differential expression results for target genes were extracted as reported, without further normalization or transformation, from “Additional file 3. Results of the proteomic analysis of AD brain microvessel extracts”.

### Surface protein expression correlation analysis

For each snRNA-seq study (except for ^37^), for each disease status and brain region group, aggregate CPM values were calculated for all genes encoding transmembrane and GPI-APs (see Supplementary Data File 1) expressed in these datasets. Pairwise Spearman’s rank correlation coefficients were then computed between these group-level aggregate CPM profiles to quantify the similarity in global surface protein transcriptomic expression between different brain regions and neurodegenerative conditions.

### Patient-level brain shuttle target analysis and statistical methods

For each scRNA-seq and snRNA-seq dataset, gene-level CPM values were calculated for BMECs from individual patients. Due to limited sample numbers in individual scRNA-seq studies, data from ^41-43^ were combined to perform a single, unified analysis. In contrast, analyses for snRNA-seq and proteomic datasets were conducted separately for each dataset. For the Erickson et al. proteomic dataset ^44^, expression values for BMVs were taken directly from the original data as protein chromatographic peak areas and further normalized, as discussed above. Transcriptomic and, when available, proteomic data for eleven brain shuttle target genes (TFRC, SLC3A2, IGF1R, TMEM30A, ALPL, SLC2A1, LRP1, LRP8, LDLR, INSR, and CA4) were extracted for downstream analysis.

Patients were stratified based on the specific research question and dataset: by brain region and disease status for Fig. 2, 3, and Supplementary Fig.1 and 2; by sex and disease status for Fig. 4 and Supplementary Fig.3; and by age bracket for Fig. 5 and Supplementary Fig. 4. All results were visualized using bar plots or violin plots to show brain shuttle target expression in relevant strata.

All statistical analyses were performed in R (v4.x) using the lme4, lmerTest, and emmeans packages for mixed-effects models, and base R for ANOVA and t-tests. For each analysis, 11 brain shuttle target genes were tested independently, and normality of residuals was assessed using the Shapiro-Wilk test. Where normality was violated on the raw scale, a log2(Expression + 1) transformation was applied if it resolved the violations; otherwise, analyses were performed on raw values. All raw p-values were corrected for multiple comparisons between genes using the Benjamini-Hochberg (BH) method to control the false discovery rate. A summary of the total number of statistical comparisons conducted for each figure and analysis, including the source studies, specific factors assessed, statistical methods used, and the number of comparisons reaching statistical significance, is provided in Supplementary Table 1. Variance decomposition was also performed for all analyses by computing the percentage of total sum of squares attributable to each factor. Results of the variance decomposition analysis are reported in Table 1, Supplementary Table 3 and Supplementary Data File 2.

For datasets in which the same patient contributed samples from multiple brain regions (Figures 2b–f, 3e/f, Supplementary Figures 1a–f, 2e/f), linear mixed-effects models were used (Expression ∼ Disease × Region + (1|PatientID)), with Disease and Region as fixed effects and PatientID as a random intercept to account for within-patient correlation. Post-hoc pairwise comparisons via estimated marginal means with Tukey adjustment were performed to assess (1) disease effects within each brain region (Control vs. AD) and (2) regional differences within each disease state.

For datasets pooling data from multiple studies (Figures 4a, 5a/b, Supplementary Figures 3a, 4a/b), linear mixed-effects models were used with the factor(s) of interest as fixed effects and study as a random intercept to absorb batch effects. Post-hoc pairwise comparisons were performed via emmeans with Tukey adjustment. For Figure 5b and Supplementary Figure 4b, log2 transformation was applied.

Two-way ANOVA (Expression ∼ Factor1 × Factor2) was used for single-study datasets with two crossed factors (Figure 4b/c and Supplementary Figure 3b/c: Disease x Sex; Figure 5c and Supplementary Figure 4c: Disease x Age Group). BH correction was applied separately for each factor’s p-values over all tested genes, and post-hoc pairwise comparisons were performed using Tukey’s HSD. Data from Erickson et al. proteomic dataset (Figures 4c, 5c and Supplementary Figures 3c, 4c) were analyzed on log2-transformed values.

For two-group comparisons of disease versus control within a single study and region (Figures 3a/b, 4d, Supplementary Figures 2a/b, 3d), Welch’s two-sample t-test was used. Data from Erickson et al. proteomic dataset (Figure 4d, 5c and Supplementary Figure 3d: Right Panel) were analyzed on log2-transformed values. For the three-group comparison (Control vs. c9ALS vs. sALS: Figures 3c/d, Supplementary Figures 2c/d), one-way ANOVA was used with Tukey’s HSD for post-hoc pairwise comparisons.

In Supplementary Table 2, differential expression results for brain shuttle targets of interest extracted directly from the supplementary material (“Additional file 3. Results of the proteomic analysis of AD BMV extracts”) of ^45^ are reported, with only the p-values being converted from log10 p-values to standard p-values. Detailed information on sample composition, demographic variables, and brain region assignments for each dataset is available in Supplementary Data File 3.

### Analysis of inter-individual variation in brain shuttle target expression

To assess inter-individual variation in brain shuttle target gene transcript and protein expression, we analyzed the healthy and/or AD groups separately within each dataset. These analyses were performed for eleven BBB-associated brain shuttle target genes (TFRC, SLC3A2, IGF1R, TMEM30A, ALPL, SLC2A1, LRP1, LRP8, LDLR, INSR, and CA4). For scRNA-seq, healthy controls ^41-43^ were merged for analysis. The snRNA-seq ^35-37^ and proteomic ^44^ datasets were analyzed independently. For each gene of interest and within each group, individuals were stratified as high or low expressors based on whether their expression value was above or below the group median. Expression range (minimum to maximum), median, fold difference between the highest and lowest expressing individuals, and mean expression of high and low expressor subgroups were calculated for each gene and reported in Table 1 and Supplementary Table 3.

Variance decomposition was performed for every statistical analysis in Figures 2-5 to quantify the proportion of total variance in gene expression attributable to each factor tested versus the residual. For all analysis types, the total sum of squares (SS_total) was computed as the sum of squared deviations from the grand mean. For Welch t-test analyses (Figs. 3a, 4d), variance decomposition was performed by fitting the equivalent one-way ANOVA (Expression ∼ Disease) and computing eta-squared as SS_between / SS_total. For one-way and two-way ANOVA analyses (Figs. 3b, 4b, 4c, 5c), the sequential (Type I) sum of squares for each factor and their interaction (where applicable) were extracted directly from the ANOVA table and expressed as a percentage of SS_total, with the residual comprising the remainder. For linear mixed-effects model analyses (Figs. 2, 3c, 4a, 5a, 5b), the sum of squares for each fixed effect was extracted from the Type III ANOVA table of the fitted model (via lmerTest), and each was expressed as a percentage of SS_total. The residual was computed as SS_total minus the sum of all fixed-effect SS; in these models, the residual therefore encompasses inter-individual biological variability, measurement noise and variance associated with the random effects. Full results of the variance decomposition analysis, reporting the contribution of each factor tested and the residual, are reported in Supplementary Data File 2. The contribution of residual variance (i.e., variance not attributable to the fixed effects tested) is also reported in Table 1.

### PMI–expression correlation analysis

For the Erickson et al. proteomic dataset ^44^ the relationship between PMI and brain shuttle target expression levels was assessed by calculating Spearman correlation coefficients between PMI and the expression value for each gene in each group, and the results are reported in Supplementary Fig. 5. All raw p-values were corrected for multiple comparisons across genes using the Benjamini-Hochberg (BH) method, and are reported on each of the plots. The aim was to determine whether PMI confounded measurements of inter-individual variation in brain shuttle target abundance. A significant positive correlation between PMI and brain shuttle target abundance would support the presence of artifactual inter-individual differences in expression resulting from differing PMIs.

## Acknowledgements

We want to thank Jana Crespo for her help with the brain drawings.

## Funding

Massachusetts Life Science Center Novel Therapeutics Delivery Grants 2021 and 2024. ISRAs: Merck, Eisai, Lilly, BMS, Lundbeck, Visterra, Alnylam

## Author contributions

Conceptualization: ARSM, SS, JG

Methodology: ARSM, SS, VC, JG, EL, RH, SH, ST, TL, IG, CS, MJ, VM

Investigation: ARSM, SS, VC, JG, EL, RH, SH, ST, TL, IG, CS, MJ, VM

Visualization: ARSM, SS, EM, RH

Funding acquisition: ARSM, JG, DI

Project administration: ARSM, JG, DI

Supervision: JG, DI

Writing – original draft: ARSM, SS, VC, JG, EL, RH, SH, ST, TL, IG, CS, MJ, VM

Writing – review & editing: ARSM, SS, VC, JG, EL, RH, SH, ST, TL, IG, CS, MJ, VM, DI

## Competing Interests

Authors declare that they have no competing interests.

## Data and Materials Availability

All data used in this study were obtained from publicly available sources, as listed in the Methods section. These datasets are accessible through GEO and dbGaP.

## List of Supplementary Materials

**Supplementary Fig. 1:**
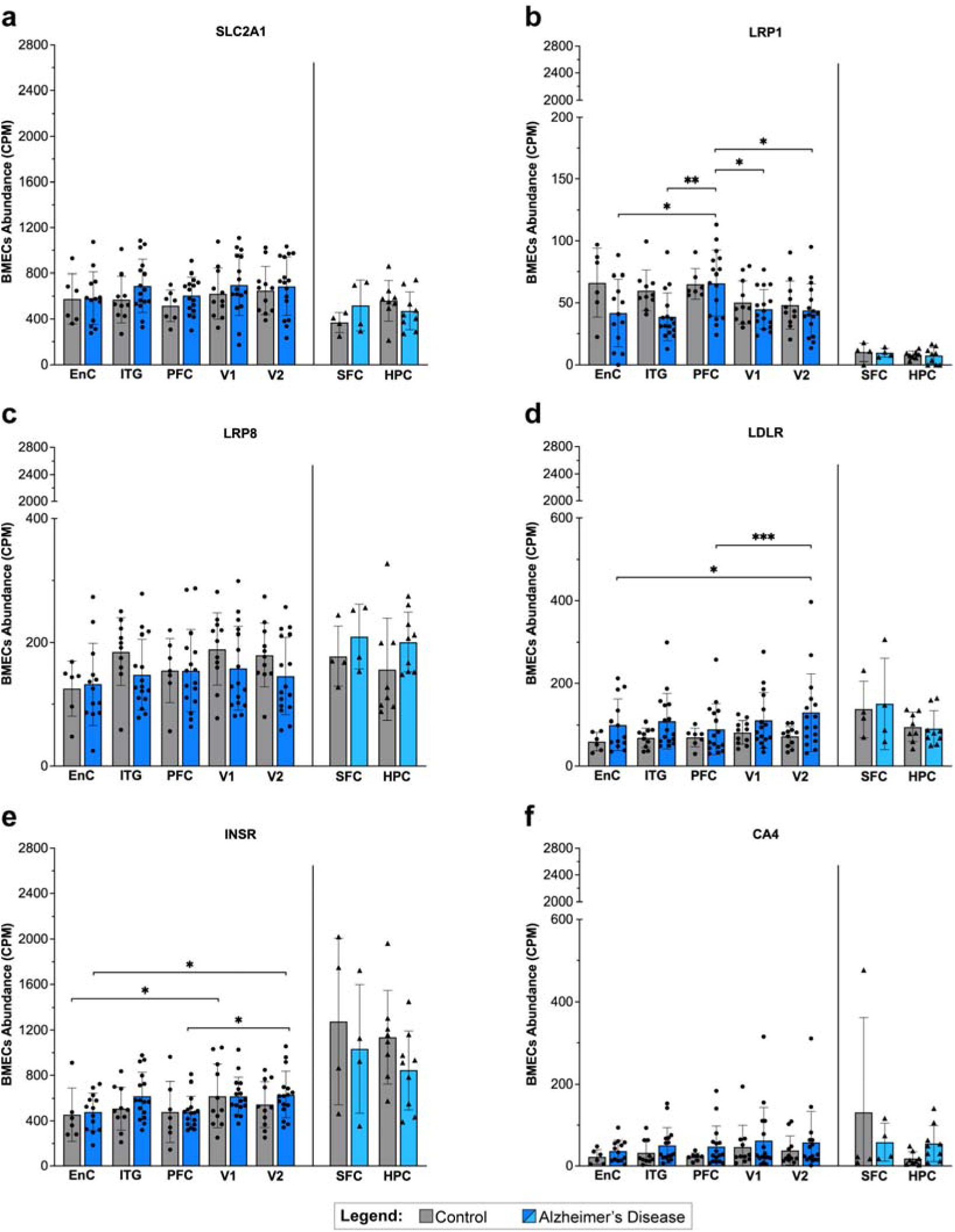
Additional brain shuttle target genes show conserved expression between brain regions and disease states in Alzheimer’s and control samples. **(a–f)** Expression levels (counts per million, CPM) of six additional brain shuttle target genes (SLC2A1, LRP1, LRP8, LDLR, INSR, and CA4) in brain microvascular endothelial cells (BMECs), measured by snRNA-seq in cognitively normal control and Alzheimer’s disease (AD) patients in seven brain regions: EC, ITG, PFC, V1, V2, SFC, and HPC. Data from ^36^ (circles) and ^35^ (triangles). Linear mixed-effects models (Disease × Region, with PatientID as random intercept) were performed within each dataset, with post-hoc pairwise comparisons via estimated marginal means with Tukey adjustment; graphs show mean ± SD, *p≤0.05, **p≤0.01, ***p≤0.001, BH-corrected over all tested genes.

**Supplementary Fig.2:**
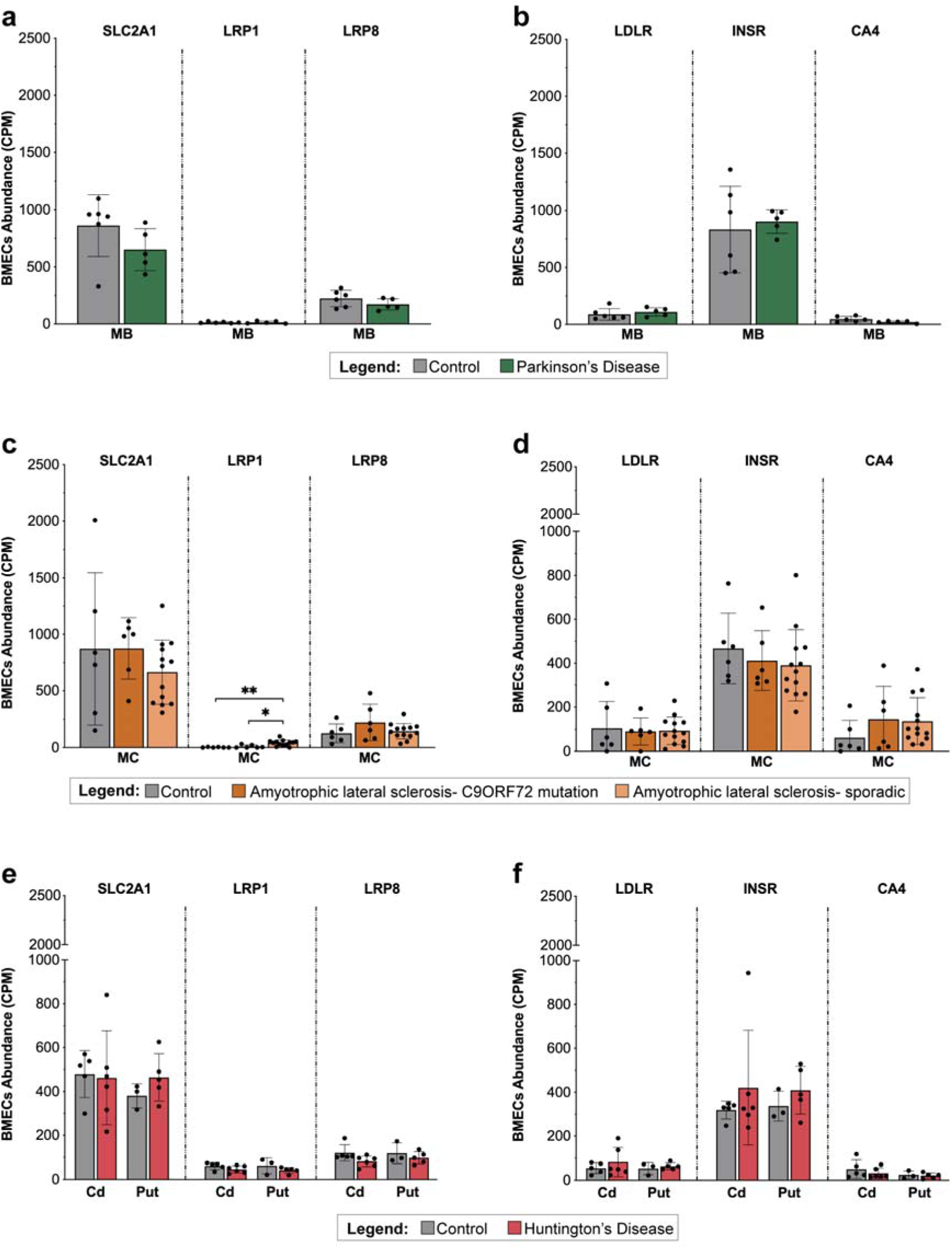
Additional brain shuttle target genes show no significant expression differences between neurodegenerative disease states and brain regions. **(a–f)** Expression (counts per million, CPM) of six additional brain shuttle target genes in brain microvascular endothelial cells (BMECs), measured by snRNA-seq between three independent neurodegenerative disease datasets. **(a, b)** Midbrain from control (grey, n = 6) and Parkinson’s disease (PD; green, n = 5) patients. **(c, d)** Motor cortex from control (grey, n = 6), C9ORF72-associated ALS (c9ALS; dark orange, n = 6), and sporadic ALS (sALS; light orange, n = 13) patients. **(e, f)** Caudate and putamen from control (Cd: n = 4, Pt: n = 3) and Huntington’s disease (HD; red; Cd: n = 6, Pt: n = 5) patients. Expression values are shown for individual patients. Welch’s t-test was used for Smajic et al. (Control vs. PD: Panels a and b); one-way ANOVA with Tukey’s HSD post-hoc test was used for Pineda et al. (Control vs. c9ALS vs. sALS: Panels c and d); and a linear mixed-effects model (Disease × Region, with PatientID as random intercept) with post-hoc pairwise comparisons via estimated marginal means with Tukey adjustment was used for Garcia et al. (Control vs. HD: Panels e and f). All p-values were BH-corrected; graphs show mean ± SD, *p≤0.05, **p≤0.01.

**Supplementary Fig. 3:**
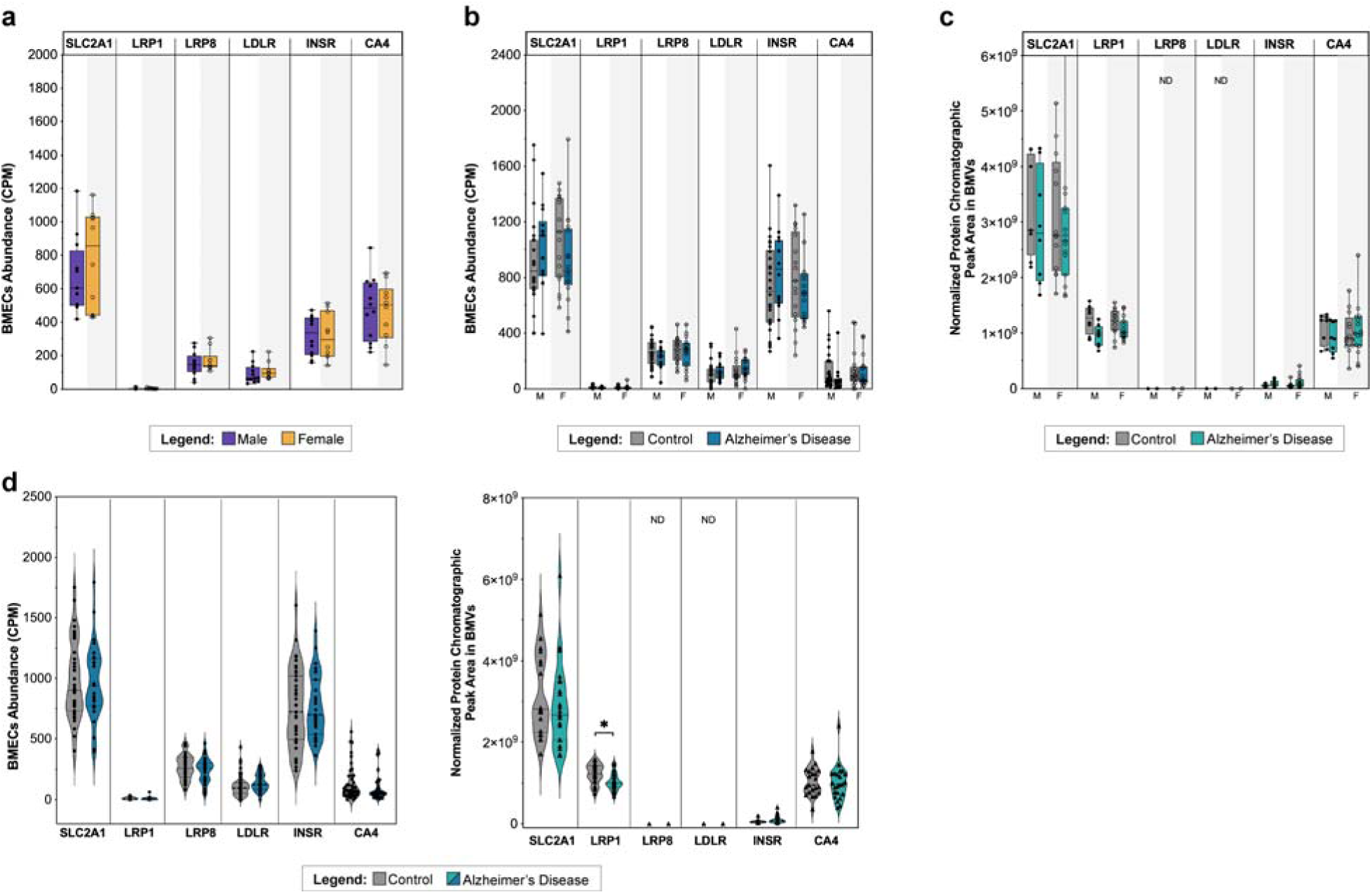
Additional brain shuttle target genes show minimal sex- or disease-related differences in AD and control samples. **(a–c)** Expression of six additional brain shuttle target genes in brain microvascular endothelial cells (BMECs) or microvessels (BMVs), stratified by sex (males: filled black circles; females: open circles) and Alzheimer’s disease (AD) status (control: grey bars, AD: blue/teal bars). **(a)** scRNA-seq data from healthy control individuals ^41-43^: males (purple, n = 12), females (yellow, n = 10). **(b)** snRNA-seq data from ^37^: control (grey) and AD (blue) individuals, control males (n = 26), AD males (n = 14), control females (n = 17), AD females (n = 17). **(c)** Proteomic data ^44^: control (grey) and AD (teal) BMVs, control males (n = 8), AD males (n = 8), control females (n = 13), AD females (n = 15). **(d)** For both snRNA-seq ^37^ and proteomic datasets ^44^, additional analyses were performed by combining males and females within control and AD groups. Graphs display individual values and meanD±DSD. A linear mixed-effects model (Expression ∼ Sex + (1|Study)) was used for the scRNA-seq data (panel a); two-way ANOVA (Disease × Sex) with Tukey’s HSD post-hoc test was used for Sun et al. and Erickson et al. sex analyses (panels b and c, respectively); and Welch’s t-test was used for Sun et al. and Erickson et al. disease-only comparisons (panel d). All p-values were BH-corrected; graphs show mean ± SD; non-detected proteins are indicated as ND. All statistical comparisons were non-significant.

**Supplementary Fig. 4:**
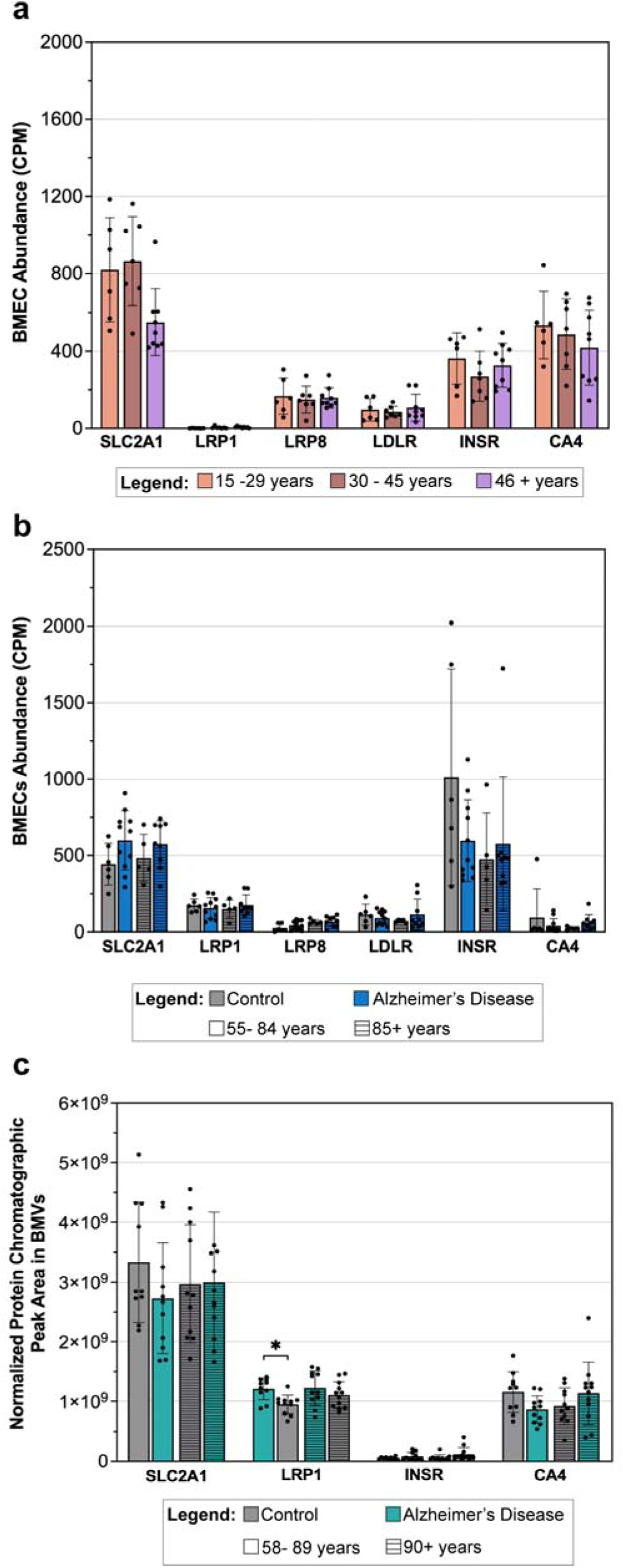
Additional brain shuttle target gene expression remains stable with age at transcript and protein levels. **(a–c)** Expression of six additional brain shuttle target genes in brain microvascular endothelial cells (BMECs) or brain microvessels (BMVs), stratified by disease status and age, using sc/snRNA-seq or proteomics. **(a)** scRNA-seq data ^41-43^ from previously described healthy control datasets, showing BMEC transcript levels (CPM) in individuals aged 15–29 (n = 6), 30–45 (n = 7), and >45 (n = 9). **(b)** snRNA-seq data from ^35, 36^, combining prefrontal (PFC) and superior frontal cortex (SFC) samples. Control (grey) and AD (blue), solid bars indicate 55–84 years and horizontally striped bars indicate ≥85 years (control: n = 6/5; AD: n = 11/9, respectively). **(c)** Proteomic data ^44^, showing normalized protein abundance in BMVs from SFC of Control (grey) and AD (teal); solid bars indicate 58–89 years and horizontally striped bars indicate ≥90 years (control: n = 10/11; AD: n = 11/12, respectively). A linear mixed-effects model (Expression ∼ AgeGroup + (1|Study)) with post-hoc pairwise comparisons via estimated marginal means with Tukey adjustment was used for the scRNA-seq data (panel a); a linear mixed-effects model (Disease × AgeGroup + (1|Study)) with Tukey-adjusted post-hoc comparisons was used for Yang/Bryant et al. (panel b); and two-way ANOVA (Disease × AgeGroup) with Tukey’s HSD post-hoc test was used for Erickson et al. (panel c). All p-values were BH-corrected; graphs show mean ± SD, *p≤0.05.

**Supplementary Fig.5:**
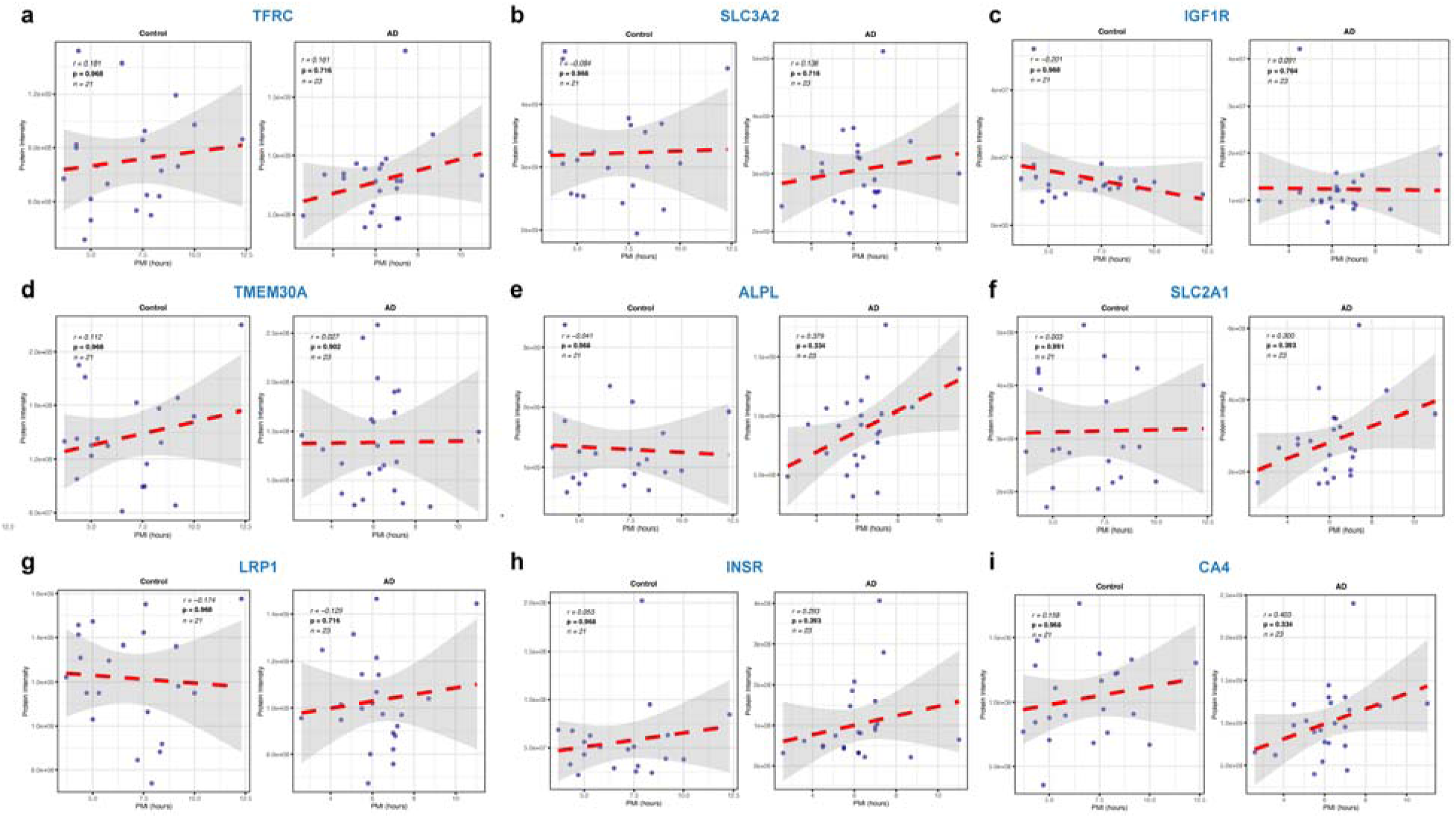
Post-mortem interval does not correlate with expression of brain shuttle target proteins in human brain microvessel data from Erickson et al ^44^. **(a–i)** Spearman correlation between post-mortem interval (PMI, hours) and abundance of brain shuttle target proteins (TFRC **(a)**, SLC3A2 **(b)**, IGF1R **(c)**, TMEM30A **(d)**, ALPL **(e)**, SLC2A1 **(f)**, LRP1 **(g)**, INSR **(h)**, CA4 **(i)**) in brain microvessels (BMVs) from control and AD subjects. Each panel displays the correlation for an individual protein in control and AD groups, with points representin individual patient samples and lines showing linear regression fit with 95% confidence intervals (gray shading). Th Spearman correlation coefficient (r), BH-corrected p-value, and sample size (n) are indicated on each plot. N significant correlations were observed for any brain shuttle target in either control or AD group.

**Supplementary Table 1:** Total number of statistical comparisons and significant results across analyses. Summary of the total number of statistical comparisons conducted for each figure and analysis, including the source studies, specific factors assessed, statistical methods used, and the number of comparisons reaching statistical significance.

| Figure(s) | Source Study for Each Figure Panel | Factors Compared | Statistical Test | Total Comparisons | Significant Results (p < 0.05) |
| --- | --- | --- | --- | --- | --- |
| Fig 2;<br>Supp Fig 1 | All Panels: Bryant et al. 2023 and Yang et al. 2022 | Disease (Control vs AD) and brain region | Mixed effects model (BH correction) and posthoc pairwise comparison (Tukey HSD correction) | 319 | 11 |
| Fig 3;<br>Supp Fig 2 | a. Smajic et al. 2022<br>b. Pineda et al. 2024<br>c. Garcia et al. 2022 | Disease (Control vs PD/ALS/HD) and brain region | a. Welch's t-test (BH correction)<br>b. One way ANOVA (BH correction) and posthoc pairwise comparison (Tukey HSD correction)<br>c. Mixed effects model (BH correction) and posthoc pairwise comparison (Tukey HSD correction) | 88 | 4 |
| Fig 4;<br>Supp Fig 3 | a. Winkler et al. 2022, Xie et al. 2024, and Wälchli et al. 2024<br>b. Sun et al. 2023<br>c. Erickson et al. 2024<br>d. Sun et al. 2023 and Erickson et al. 2024 | Disease (Control vs AD) and gender | a. Mixed effects model (BH correction)<br>b. and c. Two way ANOVA (BH correction) and posthoc pairwise comparison (Tukey HSD correction)<br>d. Welch's t-test (BH correction) | 111 | 3 |
| Fig 5;<br>Supp Fig 4 | a. Winkler et al. 2022, Xie et al. 2024, and Wälchli et al. 2024<br>b. Bryant et al. 2023 and Yang et al. 2022<br>c. Erickson et al. 2024 | Disease (Control vs AD) and age | a. Mixed effects model (BH correction) and posthoc pairwise comparison (Tukey HSD correction)<br>b. Mixed effects model (BH correction) and posthoc pairwise comparison (Tukey HSD correction)<br>c. Two way ANOVA (BH correction) and posthoc pairwise comparison (Tukey HSD correction) | 113 | 1 |

**Supplementary Table 2:**
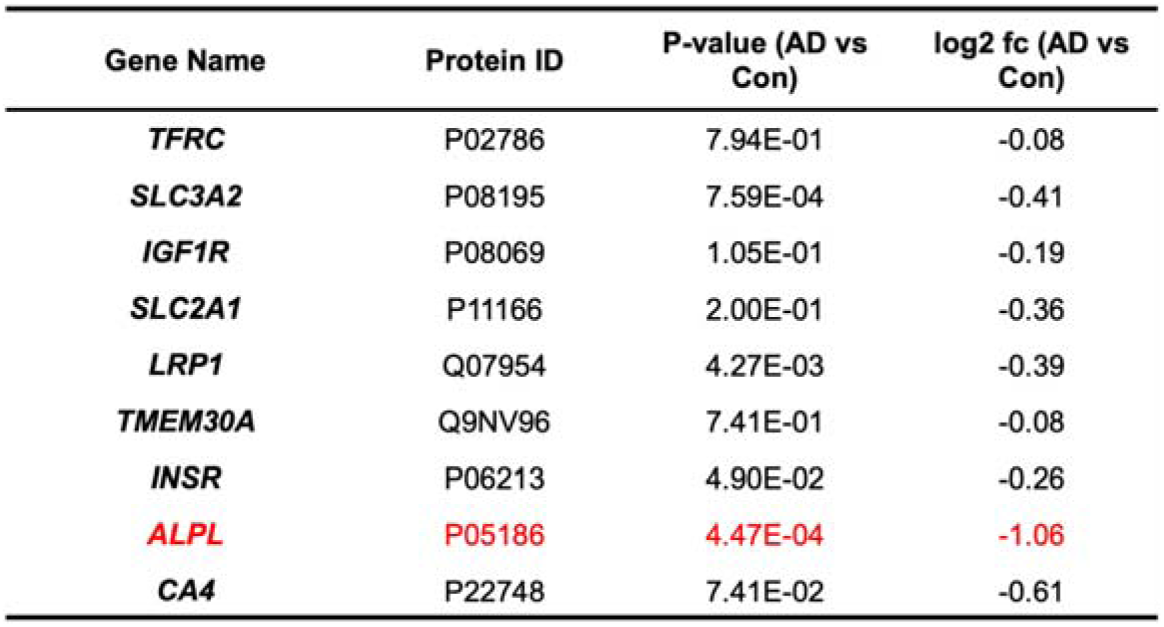
Additional proteomic analysis of differences in brain shuttle target expression between control and Alzheimer’s Disease subjects. Differential expression data for the brain shuttle targets of interest were extracted directly from the supplementary file of the original publication^45^. Among these genes, only ALPL was significantly downregulated in AD compared to control (p < 0.05, |logDFC| ≥ 1); all other brain shuttle targets did not differ.

**Supplementary Table 3:**
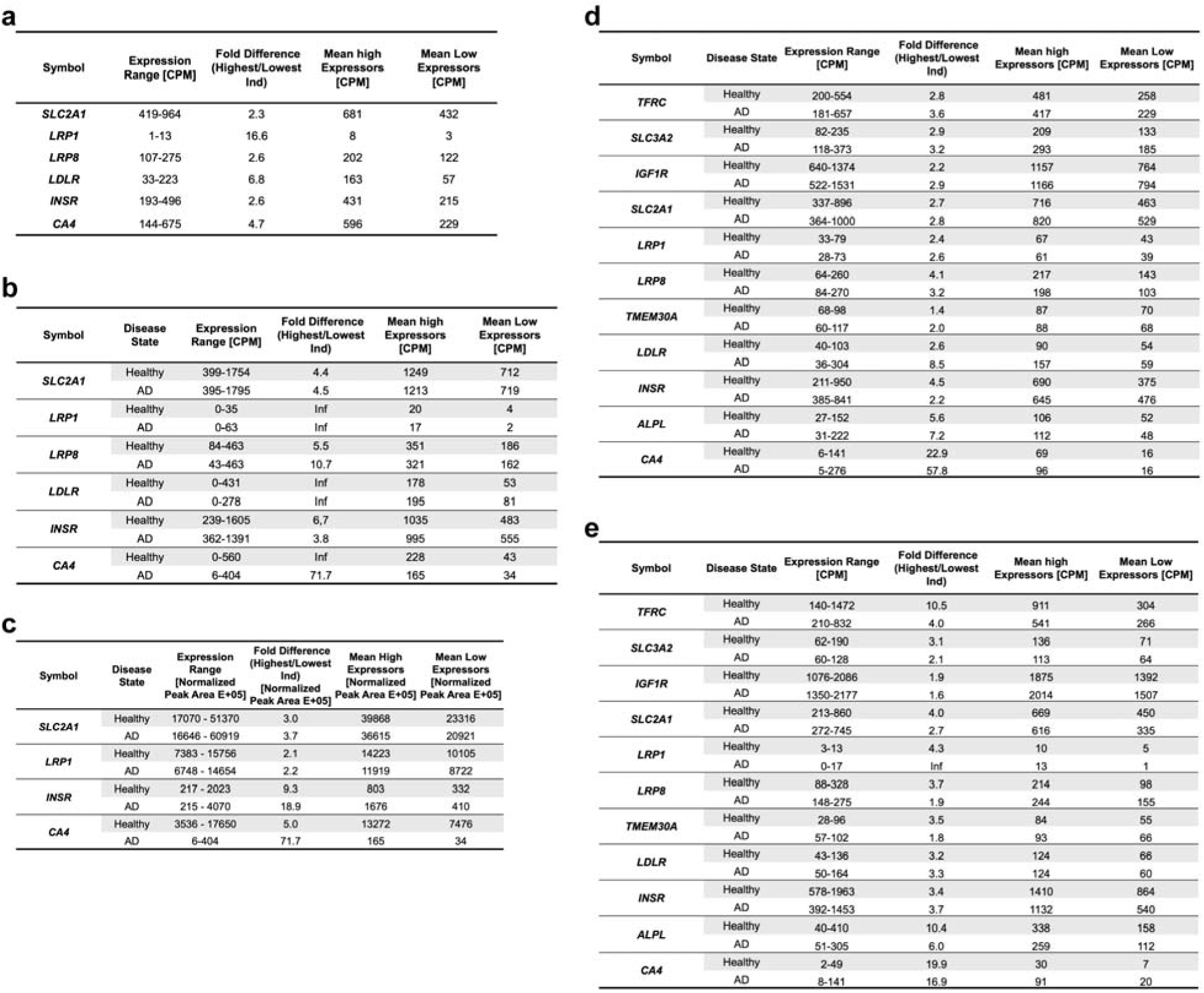
Inter-individual variation in expression of brain shuttle target genes between transcriptomic and proteomic datasets. . **(a)** scRNA-seq from healthy controls ^41-43^ (n = 22), **(b)** snRNA-seq from control (n = 43) and AD (n = 31) individuals ^37^, and **(c)** proteomic abundance from control (n = 21) and AD (n = 23) individuals^44^. **(d)** snRNAseq data^36^ (n = 6 and 13 for control and AD subjects, respectively), representing all brain regions, merged. **(e)** snRNAseq data^35^ representing hippocampus (n = 8 and 9 for control and AD subjects, respectively). Within each group, individuals were stratified as high or low expressors based on the group median. Table reports gene symbol, expression range with median, fold-difference between highest and lowest expressors, and mean expression in high vs. low expressors.

**Supplementary Data File 1:** Comprehensive list of genes encoding surface-exposed transmembrane (TM) proteins and GPI-anchored proteins (GPI-APs) used for downstream analysis. Columns: hu_uniprot, human UniProt accession ID; hu_gene, human gene symbol; ms_gene, mouse ortholog gene symbol; ms_uniprot, mouse ortholog UniProt accession ID; origin, protein classification (TM or GPI-AP).

**Supplementary Data File 2:** Full variance decomposition results for all analyses. Each tab corresponds to a main figure and its corresponding supplementary figure, with results for each panel (dataset/analysis) within. For each gene, the percentage of total variance attributable to each fixed effect, their interaction (where applicable), and the residual is reported.

**Supplementary Data File 3:** Sample composition, demographic variables, and brain regions analyzed for each dataset used in this study, derived from the supplementary materials of the original publications. Each tab corresponds to a dataset/analysis.

## Notes

### Competing Interest Statement

The authors have declared no competing interest.

### Summary of Updates

The middle name "Pato" of the first author has been removed

## References

1. Pardridge, W.M. Receptor-mediated drug delivery of bispecific therapeutic antibodies through the blood-brain barrier. Front Drug Deliv 3 (2023).

2. Banks, W.A. From blood-brain barrier to blood-brain interface: new opportunities for CNS drug delivery. Nat Rev Drug Discov 15, 275–292 (2016).

3. Pulgar, V.M. Transcytosis to Cross the Blood Brain Barrier, New Advancements and Challenges. Front Neurosci 12, 1019 (2018).

4. Abbott, N.J., Patabendige, A.A., Dolman, D.E., Yusof, S.R. & Begley, D.J. Structure and function of the blood-brain barrier. Neurobiol Dis 37, 13–25 (2010).

5. Barker, S.J. et al. Targeting the transferrin receptor to transport antisense oligonucleotides across the mammalian blood-brain barrier. Sci Transl Med 16, eadi2245 (2024).

6. Niewoehner, J. et al. Increased brain penetration and potency of a therapeutic antibody using a monovalent molecular shuttle. Neuron 81, 49–60 (2014).

7. Molino, Y. et al. Use of LDL receptor-targeting peptide vectors for in vitro and in vivo cargo transport across the blood-brain barrier. FASEB J 31, 1807–1827 (2017).

8. P, X. et al. The role of LRP1 in Abeta efflux transport across the blood-brain barrier and cognitive dysfunction in diabetes mellitus. Neurochem Int 160, 105417 (2022).

9. Sinha, R.K. et al. Apolipoprotein E Receptor 2 Mediates Activated Protein C-Induced Endothelial Akt Activation and Endothelial Barrier Stabilization. Arterioscler Thromb Vasc Biol 36, 518–524 (2016).

10. Boado, R.J., Lu, J.Z., Hui, E.K., Sumbria, R.K. & Pardridge, W.M. Pharmacokinetics and brain uptake in the rhesus monkey of a fusion protein of arylsulfatase a and a monoclonal antibody against the human insulin receptor. Biotechnol Bioeng 110, 1456–1465 (2013).

11. Shin, J.W. et al. Grabody B, an IGF1 receptor-based shuttle, mediates efficient delivery of biologics across the blood-brain barrier. Cell Rep Methods 2, 100338 (2022).

12. Veys, K. et al. Role of the GLUT1 Glucose Transporter in Postnatal CNS Angiogenesis and Blood-Brain Barrier Integrity. Circ Res 127, 466–482 (2020).

13. Khoury, N. et al. Fc-engineered large molecules targeting blood-brain barrier transferrin receptor and CD98hc have distinct central nervous system and peripheral biodistribution. Nat Commun 16, 1822 (2025).

14. Moyer, T.C. et al. Highly conserved brain vascular receptor ALPL mediates transport of engineered AAV vectors across the blood-brain barrier. Mol Ther 33, 3902–3916 (2025).

15. Shay, T.F. et al. Primate-conserved carbonic anhydrase IV and murine-restricted LY6C1 enable blood-brain barrier crossing by engineered viral vectors. Sci Adv 9, eadg6618 (2023).

16. Lessard, E. et al. Pharmacokinetics and Pharmacodynamic Effect of a Blood-Brain Barrier-Crossing Fusion Protein Therapeutic for Alzheimer’s Disease in Rat and Dog. Pharm Res 39, 1497–1507 (2022).

17. Sonoda, H. et al. A Blood-Brain-Barrier-Penetrating Anti-human Transferrin Receptor Antibody Fusion Protein for Neuronopathic Mucopolysaccharidosis II. Mol Ther 26, 1366–1374 (2018).

18. Denali’s blood-brain barrier shuttle shows promise. C&EN Global Enterprise 99, 15-15 (2021).

19. Grimm, H.P. et al. Delivery of the Brainshuttle amyloid-beta antibody fusion trontinemab to non-human primate brain and projected efficacious dose regimens in humans. MAbs 15, 2261509 (2023).

20. Pizzo, M.E. et al. Transferrin receptor-targeted anti-amyloid antibody enhances brain delivery and mitigates ARIA. Science 389, eads3204 (2025).

21. Okuyama, T. et al. A Phase 2/3 Trial of Pabinafusp Alfa, IDS Fused with Anti-Human Transferrin Receptor Antibody, Targeting Neurodegeneration in MPS-II. Mol Ther 29, 671–679 (2021).

22. Giugliani, R. et al. Iduronate-2-sulfatase fused with anti-hTfR antibody, pabinafusp alfa, for MPS-II: A phase 2 trial in Brazil. Mol Ther 29, 2378–2386 (2021).

23. An, S. et al. A brain-shuttled antibody targeting alpha synuclein aggregates for the treatment of synucleinopathies. NPJ Parkinsons Dis 11, 254 (2025).

24. Alfaidi, M., Barker, R.A. & Kuan, W.L. An update on immune-based alpha-synuclein trials in Parkinson’s disease. J Neurol 272, 21 (2024).

25. Shinohara, M., Tachibana, M., Kanekiyo, T. & Bu, G. Role of LRP1 in the pathogenesis of Alzheimer’s disease: evidence from clinical and preclinical studies. J Lipid Res 58, 1267–1281 (2017).

26. Winkler, E.A. et al. GLUT1 reductions exacerbate Alzheimer’s disease vasculo-neuronal dysfunction and degeneration. Nat Neurosci 18, 521–530 (2015).

27. Leclerc, M. et al. Lower GLUT1 and unchanged MCT1 in Alzheimer’s disease cerebrovasculature. J Cereb Blood Flow Metab 44, 1417–1432 (2024).

28. Petralla, S. et al. Increased Expression of Transferrin Receptor 1 in the Brain Cortex of 5xFAD Mouse Model of Alzheimer’s Disease Is Associated with Activation of HIF-1 Signaling Pathway. Mol Neurobiol 61, 6383–6394 (2024).

29. Lewitt, M.S. & Boyd, G.W. Role of the Insulin-like Growth Factor System in Neurodegenerative Disease. Int J Mol Sci 25 (2024).

30. Yang, A.C. et al. Physiological blood-brain transport is impaired with age by a shift in transcytosis. Nature 583, 425–430 (2020).

31. Torres, V.O. et al. Transferrin receptor-mediated transport at the blood-brain barrier is elevated during early development and maintained across aging and in an Alzheimer’s mouse model. J Cereb Blood Flow Metab, 271678X251361997 (2025).

32. Zhang, W. et al. Differential expression of receptors mediating receptor-mediated transcytosis (RMT) in brain microvessels, brain parenchyma and peripheral tissues of the mouse and the human. Fluids Barriers CNS 17, 47 (2020).

33. Sweeney, M.D., Sagare, A.P. & Zlokovic, B.V. Blood-brain barrier breakdown in Alzheimer disease and other neurodegenerative disorders. Nat Rev Neurol 14, 133–150 (2018).

34. Knox, E.G., Aburto, M.R., Clarke, G., Cryan, J.F. & O’Driscoll, C.M. The blood-brain barrier in aging and neurodegeneration. Mol Psychiatry 27, 2659–2673 (2022).

35. Yang, A.C. et al. A human brain vascular atlas reveals diverse mediators of Alzheimer’s risk. Nature 603, 885–892 (2022).

36. Bryant, A. et al. Endothelial Cells Are Heterogeneous in Different Brain Regions and Are Dramatically Altered in Alzheimer’s Disease. J Neurosci 43, 4541–4557 (2023).

37. Sun, N. et al. Single-nucleus multiregion transcriptomic analysis of brain vasculature in Alzheimer’s disease. Nat Neurosci 26, 970–982 (2023).

38. Smajic, S. et al. Single-cell sequencing of human midbrain reveals glial activation and a Parkinson-specific neuronal state. Brain 145, 964–978 (2022).

39. Garcia, F.J. et al. Single-cell dissection of the human brain vasculature. Nature 603, 893–899 (2022).

40. Pineda, S.S. et al. Single-cell dissection of the human motor and prefrontal cortices in ALS and FTLD. Cell 187, 1971–1989 e1916 (2024).

41. Winkler, E.A. et al. A single-cell atlas of the normal and malformed human brain vasculature. Science 375, eabi7377 (2022).

42. Xie, Y. et al. Single-cell dissection of the human blood-brain barrier and glioma blood-tumor barrier. Neuron 112, 3089–3105 e3087 (2024).

43. Walchli, T. et al. Single-cell atlas of the human brain vasculature across development, adulthood and disease. Nature 632, 603–613 (2024).

44. Erickson, M.A. et al. Data-independent acquisition proteomic analysis of the brain microvasculature in Alzheimer’s disease identifies major pathways of dysfunction and upregulation of cytoprotective responses. Fluids Barriers CNS 21, 84 (2024).

45. Zellner, A. et al. Proteomic profiling in cerebral amyloid angiopathy reveals an overlap with CADASIL highlighting accumulation of HTRA1 and its substrates. Acta Neuropathol Commun 10, 6 (2022).

46. Wan, Y.W. et al. Meta-Analysis of the Alzheimer’s Disease Human Brain Transcriptome and Functional Dissection in Mouse Models. Cell Rep 32, 107908 (2020).

47. Gupta, A.K. et al. Comprehensive characterization of the RNA editing landscape in the human aging brains with Alzheimer’s disease. Alzheimers Dement 21, e70452 (2025).

48. Xu, J. et al. Regional protein expression in human Alzheimer’s brain correlates with disease severity. Commun Biol 2, 43 (2019).

49. Zhu, B. et al. Single-cell transcriptomic and proteomic analysis of Parkinson’s disease brains. Sci Transl Med 16, eabo1997 (2024).

50. Keo, A. et al. Transcriptomic signatures of brain regional vulnerability to Parkinson’s disease. Commun Biol 3, 101 (2020).

51. Cappelletti, C. et al. Transcriptomic profiling of Parkinson’s disease brains reveals disease stage specific gene expression changes. Acta Neuropathol 146, 227–244 (2023).

52. Esteves, A.R. & Cardoso, S.M. Differential protein expression in diverse brain areas of Parkinson’s and Alzheimer’s disease patients. Sci Rep 10, 13149 (2020).

53. Blumenreich, S. et al. Large-scale proteomics analysis of five brain regions from Parkinson’s disease patients with a GBA1 mutation. NPJ Parkinsons Dis 10, 33 (2024).

54. Zhang, X. et al. Region-specific protein abundance changes in the brain of MPTP-induced Parkinson’s disease mouse model. J Proteome Res 9, 1496–1509 (2010).

55. Bostrand, S.M.K. et al. Mapping the glial transcriptome in Huntington’s disease using snRNAseq: selective disruption of glial signatures across brain regions. Acta Neuropathol Commun 12, 165 (2024).

56. Mullari, M. et al. Characterising the RNA-binding protein atlas of the mammalian brain uncovers RBM5 misregulation in mouse models of Huntington’s disease. Nat Commun 14, 4348 (2023).

57. Grima, N. et al. Multi-region brain transcriptomic analysis of amyotrophic lateral sclerosis reveals widespread RNA alterations and substantial cerebellum involvement. Mol Neurodegener 20, 40 (2025).

58. Lee, A., Henderson, R., Arachchige, B.J., Robertson, T. & McCombe, P.A. Proteomic investigation of ALS motor cortex identifies known and novel pathogenetic mechanisms. J Neurol Sci 452, 120753 (2023).

59. Blanchette, M. et al. Regional heterogeneity of the blood-brain barrier. Nat Commun 16, 7332 (2025).

60. Van Norman, G.A. Limitations of Animal Studies for Predicting Toxicity in Clinical Trials: Is it Time to Rethink Our Current Approach? JACC Basic Transl Sci 4, 845–854 (2019).

61. Atkins, J.T. et al. Pre-clinical animal models are poor predictors of human toxicities in phase 1 oncology clinical trials. Br J Cancer 123, 1496–1501 (2020).

62. Marshall, L.J., Bailey, J., Cassotta, M., Herrmann, K. & Pistollato, F. Poor Translatability of Biomedical Research Using Animals - A Narrative Review. Altern Lab Anim 51, 102–135 (2023).

63. Ward, R.J., Zucca, F.A., Duyn, J.H., Crichton, R.R. & Zecca, L. The role of iron in brain ageing and neurodegenerative disorders. Lancet Neurol 13, 1045–1060 (2014).

64. Berg, D. et al. Brain iron pathways and their relevance to Parkinson’s disease. J Neurochem 79, 225–236 (2001).

65. Talbot, K. et al. Demonstrated brain insulin resistance in Alzheimer’s disease patients is associated with IGF-1 resistance, IRS-1 dysregulation, and cognitive decline. J Clin Invest 122, 1316-1338 (2012).

66. Moloney, A.M. et al. Defects in IGF-1 receptor, insulin receptor and IRS-1/2 in Alzheimer’s disease indicate possible resistance to IGF-1 and insulin signalling. Neurobiol Aging 31, 224–243 (2010).

67. Takasugi, N. et al. TMEM30A is a candidate interacting partner for the beta-carboxyl-terminal fragment of amyloid-beta precursor protein in endosomes. PLoS One 13, e0200988 (2018).

68. Vardy, E.R., Kellett, K.A., Cocklin, S.L. & Hooper, N.M. Alkaline phosphatase is increased in both brain and plasma in Alzheimer’s disease. Neurodegener Dis 9, 31–37 (2012).

69. Aronica, E. et al. Molecular classification of amyotrophic lateral sclerosis by unsupervised clustering of gene expression in motor cortex. Neurobiol Dis 74, 359–376 (2015).

70. Labadorf, A. et al. RNA Sequence Analysis of Human Huntington Disease Brain Reveals an Extensive Increase in Inflammatory and Developmental Gene Expression. PLoS One 10, e0143563 (2015).

71. Christodoulou, C.C., Onisiforou, A., Zanos, P. & Papanicolaou, E.Z. Unraveling the transcriptomic signatures of Parkinson’s disease and major depression using single-cell and bulk data. Front Aging Neurosci 15, 1273855 (2023).

72. Mi, X., Ye, Z.L., Zhang, X.J., Chen, X.C. & Dai, X.M. Sex- and age- differences in the expression of critical blood-brain barrier regulators: a physiological context. Biol Sex Differ 16, 67 (2025).

73. Thaysen, M. et al. Evaluating sex as a biological variable in in vitro blood-brain barrier models: insights from primary mouse brain endothelial cells. Fluids Barriers CNS (2026).

74. Weber, C.M. & Clyne, A.M. Sex differences in the blood-brain barrier and neurodegenerative diseases. APL Bioeng 5, 011509 (2021).

75. Torres, V.O. et al. Transferrin receptor-mediated transport at the blood-brain barrier is elevated during early development and maintained across aging and in an Alzheimer’s mouse model. J Cereb Blood Flow Metab 46, 20–35 (2026).

76. Faresjo, R., Sehlin, D. & Syvanen, S. Age, dose, and binding to TfR on blood cells influence brain delivery of a TfR-transported antibody. Fluids Barriers CNS 20, 34 (2023).

77. Zhou, X., et al. Proteomic Profiling Reveals Age-Related Changes in Transporter Proteins in the Human Blood-Brain Barrier. bioRxiv (2024).

78. Labots, G., Jones, A., de Visser, S.J., Rissmann, R. & Burggraaf, J. Gender differences in clinical registration trials: is there a real problem? Br J Clin Pharmacol 84, 700–707 (2018).

79. Bierer, B.E., Meloney, L.G., Ahmed, H.R. & White, S.A. Advancing the inclusion of underrepresented women in clinical research. Cell Rep Med 3, 100553 (2022).

80. Clemmensen, F.K. et al. Short-term variability of Alzheimer’s disease plasma biomarkers in a mixed memory clinic cohort. Alzheimers Res Ther 17, 26 (2025).

81. Patel, R. et al. Inter- and intra-individual variation in brain structural-cognition relationships in aging. Neuroimage 257, 119254 (2022).

82. Kim, B.H. et al. Second-generation anti-amyloid monoclonal antibodies for Alzheimer’s disease: current landscape and future perspectives. Transl Neurodegener 14, 6 (2025).

83. Robbins, C.J., Bates, K.M. & Rimm, D.L. HER2 testing: evolution and update for a companion diagnostic assay. Nat Rev Clin Oncol 22, 408–423 (2025).

84. Jorgensen, J.T. et al. A Companion Diagnostic With Significant Clinical Impact in Treatment of Breast and Gastric Cancer. Front Oncol 11, 676939 (2021).

85. Cox, J. & Mann, M. Quantitative, high-resolution proteomics for data-driven systems biology. Annu Rev Biochem 80, 273–299 (2011).

86. Liu, Y., Beyer, A. & Aebersold, R. On the Dependency of Cellular Protein Levels on mRNA Abundance. Cell 165, 535–550 (2016).

87. Vogel, C. & Marcotte, E.M. Insights into the regulation of protein abundance from proteomic and transcriptomic analyses. Nat Rev Genet 13, 227–232 (2012).

88. Miao, Z. et al. Putative cell type discovery from single-cell gene expression data. Nat Methods 17, 621–628 (2020).

